# Somatic mutations reveal the ontogeny of human microglia

**DOI:** 10.64898/2026.05.19.726366

**Authors:** Julia A. Belk, Yaowen Zhang, Quanming Shi, Lisa Ma, Raja Kalluru, Alejandro Medina Enciso, Emily E. Reilly, Jacob Weiss, Rui Li, Anna E. Eastman, Nicole Womack-Gambrel, Debasmita Paul, Arnav Chakravarthy, Syed Bukhari, Dipabarna Bhattacharya, Suyash Raj, Daniel Richard, Simone Brioschi, Matthew R. Chrostek, Daniel C. Nachun, Christopher M. Arends, Jayakrishnan Gopakumar, Isak W. Tengesdal, Ademar Bynum, Shaneice Mitchell, Katalin Sandor, Badri N. Vardarajan, Inma Cobos, Donald E. Born, Anne Brunet, Marco Colonna, Hannes Vogel, Thomas J. Montine, Jody E. Hooper, Irving L. Weissman, C. Dirk Keene, Howard Y. Chang, Siddhartha Jaiswal

**Affiliations:** Department of Pathology, Stanford University, Stanford, CA 94305, USA; Department of Dermatology, Stanford University, Stanford, CA 94305, USA; Amgen Research, South San Francisco, CA 94080, USA; Sarafan ChEM-H, Stanford University, Stanford, CA 94305, USA; Institute for Stem Cell Biology and Regenerative Medicine, Stanford University School of Medicine, Stanford, CA 94305, USA; Department of Genetics, Stanford University, Stanford, CA 94305, USA; Department of Pathology and Immunology, Washington University School of Medicine in Saint Louis, Saint Louis, MO 63110, USA; Taub Institute for Research on Alzheimer’s Disease and the Aging Brain, College of Physicians and Surgeons, Columbia University, New York, NY 10032, USA; Department of Neurology, College of Physicians and Surgeons, Columbia University and the New York Presbyterian Hospital, New York, NY 10032, USA; Department of Laboratory Medicine and Pathology, University of Washington School of Medicine, Seattle, WA 98195, USA

## Abstract

Microglia are the resident hematopoietic cells of the central nervous system^1^. In mice, microglia seed the brain during embryogenesis and can be maintained throughout life with minimal input from adult hematopoiesis^2–4^. The origins of human microglia are less clear, but recent evidence suggests that marrow-derived cells may be able to supplement the human microglial pool in certain individuals^5,6^. Here, to investigate the ontogeny of human microglia, we develop a method that uses the collection of accumulated somatic mutations which uniquely labels each clone of cells to track the infiltration of marrow-derived cells into the human brain. Applying this method to 20 aged individuals, we find evidence of an influx of marrow-derived cells into the brain in all examined individuals. Single cell analysis, including single cell lineage tracing using mitochondrial DNA variants, demonstrates that these infiltrating cells are nearly identical to microglia and can comprise a large fraction of the microglial pool. Analysis of large-scale sequencing cohorts demonstrates a protective association between most types of clonal hematopoiesis and Alzheimer’s disease. In sum, this work uncovers a widespread influx of myeloid cells into the healthy human brain which serves to reinforce the pool of human microglia and becomes common with aging.

## Main

Adult humans contain a polyclonal mixture of approximately 100,000 hematopoietic stem cells (HSCs)^7^. During aging, individual HSCs accumulate somatic alterations that can confer a fitness advantage, leading to clonal expansion^8–10^. Collectively, these expanded HSC clones yield an oligoclonal mixture in the blood and bone marrow of aged humans, termed “clonal hematopoiesis”. Clonal hematopoiesis (CH) has been associated with diverse human diseases in human cohort data, some of which have been causally validated in disease models^11–16^. For example, recent evidence suggests that a subset of CH caused by specific driver mutations (e.g., *TET2* mutations) has a protective association with Alzheimer’s disease (AD)^5,16,17^, in part due to an enhanced ability of these mutant cells to infiltrate the brain and supplement the pool of microglia^5,16^, the resident hematopoietic cells of the central nervous system.

While the origins and clonal dynamics of human blood throughout aging are largely known, the origins of human microglia are less clear. In mice, microglia develop from an erythro-myeloid progenitor which arises in the yolk sac before embryonic day 8 and seeds the brain during embryogenesis^2,18^. Transcriptomic studies support a yolk sac origin for human microglia in early development^19,20^. However, in human aging, emerging evidence suggests that HSC-derived clones with leukemia-associated mutations – such as mutations in *DNMT3A*, *TET2*, and *ASXL1* – can supplement the pool of brain-resident microglia^5,6^. These findings are unexpected because the pool of murine microglia is maintained throughout the lifespan via local self-renewal with minimal contribution from the periphery^3^. In mice, while various insults can cause the recruitment of additional myeloid cells into the brain, these cells do not fully adopt the microglial program, even after extended periods of brain residence^21–24^. It remains unclear whether the ability of peripheral hematopoietic cells to supplement the pool of human microglia is restricted to clones caused by specific driver events or instead might be a common feature of human aging. In particular, the types of clones that can infiltrate into the human brain, the extent of infiltration, the phenotype of these cells, and their potential functional role in neurodegenerative disease are not known.

### Development of PACT

Addressing these questions requires the ability to track cell clones originating in the bone marrow and evaluate the presence of these clones in the human brain. All cells acquire DNA mutations over time, which can be used as lineage tracing tools^7,25–29^. Clonal expansions become ubiquitous during aging^8–11,30–32^, and when a particular cell starts expanding, the accumulated nuclear somatic mutations present in that cell are passed on to all progeny (**Figure 1a**). When a clone is sufficiently expanded, these “passenger” mutations become detectable in low-depth sequencing data from bulk DNA. We hypothesized that the aggregated set of passenger mutations identified from clones in blood could be assessed in matched post-mortem brain specimens to evaluate the presence or absence of infiltrating cells originating in the bone marrow, an approach we refer to as Passenger Assisted Clone Tracking (PACT; **Figure 1b**).

**Figure 1:**
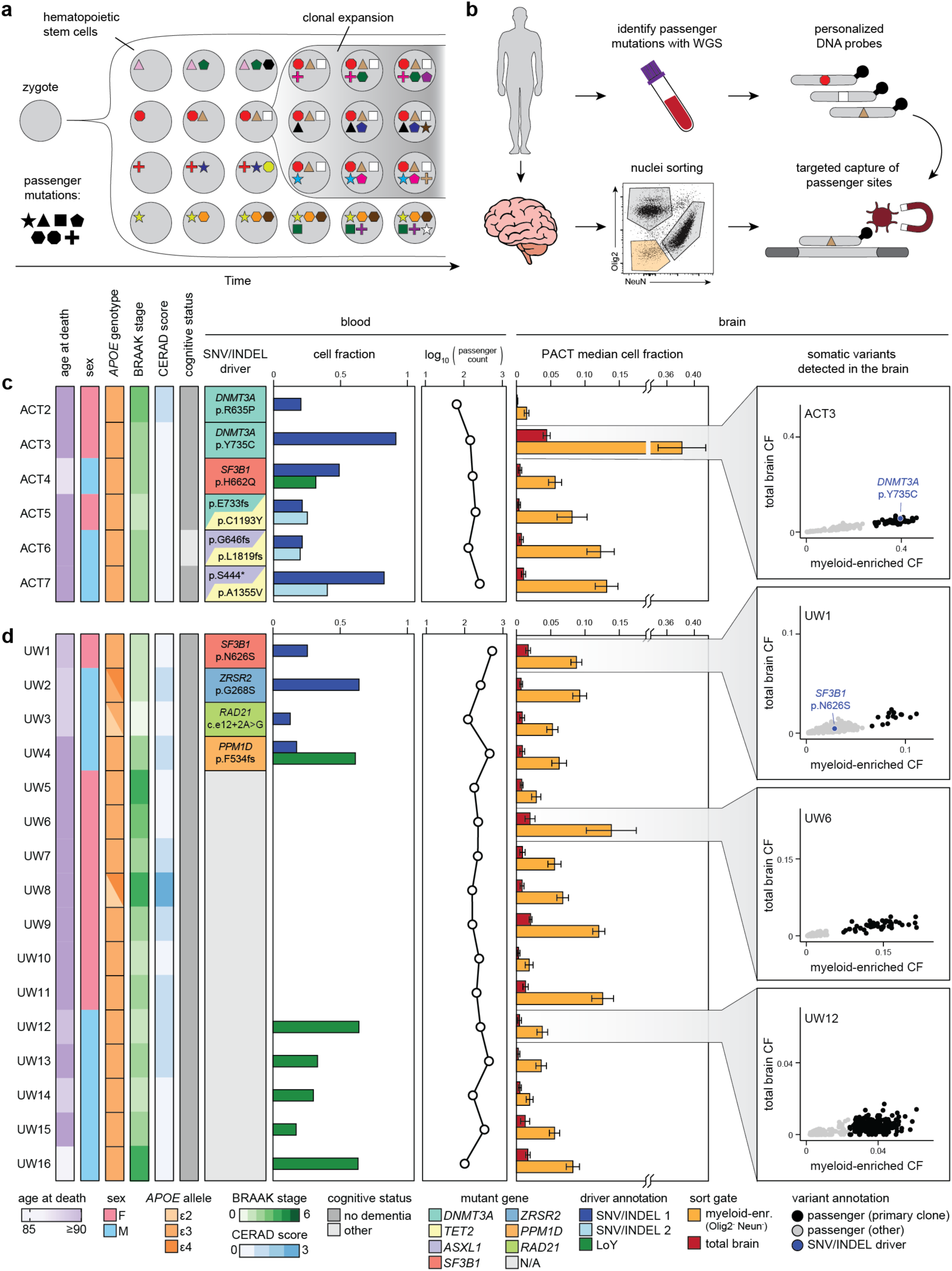
PACT reveals an influx of marrow-derived cells into the human brain. **(a)** Illustration of passenger mutation acquisition during aging. **(b)** Schematic of PACT workflow in postmortem human brain specimens. **(c)** Development of PACT using tissue specimens from donors with known hematopoietic infiltration in the brain^5^. **(d)** Application of PACT to tissue specimens from additional human donors. **(c-d)** Clinical and neuropathological characteristics (left), blood characteristics (center) and PACT results from postmortem brain specimens (right) from human donors. Error bars represent median ± median absolute deviation. WGS, whole genome sequencing; CF, cell fraction; SNV/INDEL, single nucleotide variant or insertion or deletion; LoY, loss of the Y chromosome.

To develop PACT, we first investigated individuals with CH due to known single nucleotide variant, insertion, or deletion (SNV or INDEL) driver mutations who had previously been shown to have the same driver mutations detectable in their brain (**Figure 1c**; **Supplementary Tables 1-2**)^5^. These individuals have available postmortem brain samples available from the University of Washington BioRepository and Integrated Neuropathology (BRaIN) laboratory and antemortem blood whole genome sequencing data collected as part of the AD Sequencing Project (ADSP). We identified putative passenger mutations from the blood WGS data of each individual and synthesized custom oligonucleotide probes targeting each mutation site, as well as the known driver mutation(s) (**Extended Data Figure 1a-b**). From post-mortem brain specimens, we used flow cytometry to isolate total nuclei, myeloid-enriched nuclei (NeuN^−^Olig2^−^), neurons (NeuN^+^), and oligodendrocytes (Olig2^+^; **Extended Data Figure 1c**). We extracted DNA and performed targeted capture of the personalized mutation sites for each individual in each sorted brain fraction. After sequencing, we quantified the allele frequency of each mutation in each sample.

As expected, the CH driver mutation was enriched in the myeloid-enriched fraction of the brain relative to total brain nuclei, confirming that peripherally derived cells had engrafted in the brain. We then examined the passenger mutations for each individual. We filtered the passenger mutations by requiring at least two-fold enrichment in the myeloid-enriched fraction of the brain relative to the total brain nuclei and then visualized the cell fraction of each mutation in the myeloid-enriched fraction of the brain, the total brain nuclei, and the blood (**Extended Data Figure 2a-b**). We found that the cell fraction distribution of the passenger mutations typically had at least one clear mode and we refer to this main cluster of passengers as the “primary clone.” The cell fraction of the driver mutation was well approximated by the median cell fraction of the passengers present in the primary clone, enabling us to estimate the clone size of the primary clone directly from the distribution of passenger mutations (**Extended Data Figure 2**). In certain subjects, additional clusters of passengers were present with lower cell fractions which may represent smaller clones. Finally, we compared the myeloid-enriched passenger mutations, nonspecific (likely germline) mutations, and putative variants that were not detected in the brain in order to discern features of informative variants (**Extended Data Figure 3**). These analyses revealed the most useful variant calling parameters and indicated that transitions were enriched for genuine mutations (**Extended Data Figure 3**). Together, these results demonstrate that PACT is an effective strategy to assess infiltration of HSC-derived clones into the human brain and can provide quantitative estimates of clone size, even in the absence of knowledge of the driver event(s).

### Marrow-derived cells colonize the human brain

We next sought to apply PACT to individuals without a defined SNV or INDEL driver mutation to better understand the extent to which different types of clones can infiltrate the human brain. We analyzed the blood WGS data from all individuals with postmortem brain samples available in the BRaIN laboratory, identifying driver SNVs, INDELs, and mosaic chromosomal alterations (mCAs; including loss of the Y chromosome (LoY)), and putative passenger mutations for each individual. We used curated whitelists for each type of driver event (SNV, INDEL, or mCA) to identify individuals with clonal expansions with known drivers and used the aggregate passenger burden to identify individuals with CH without known drivers^32^. We prioritized 16 individuals who had a diverse group of clonal characteristics, including unknown drivers and mCAs, for further investigation (**Figure 1d**; **Supplemental Table 2**).

PACT revealed an infiltration of clonally expanded blood cells into the brain in each individual (**Figure 1d**). As before, examining the empirical passenger distributions of each individual revealed a primary clone as well as smaller clones in some individuals (**Extended Data Figure 4**). In individuals with multiple co-existing clones, these empirical distributions also suggested clonal competition dynamics within the brain. For example, UW1 harbored an *SF3B1* mutation which was detectable in the brain but at lower cell fraction than a second clone. The second UW1 clone harbored fewer passenger mutations, suggesting that it originated earlier in life than the *SF3B1*-mutant clone (**Extended Data Figure 4**). Secondly, UW3 harbored a *RAD21* driver variant in the blood, which was not detectable in the brain, but a second unknown-driver clone was present in the brain of UW3. UW3 thus presents an example of one clone outcompeting another for brain residence (**Extended Data Figure 4**).

To better understand the passenger distributions that could result from different clonal competition scenarios within the brain, we performed simulation studies (**Extended Data Figure 5**). Because mutations accumulate with time, the number of passenger variants marking a particular clone indicates the age of the clone. Our simulations considered two co-existing hematopoietic clones of different ages, took as input the cell fraction of each clone in the blood and the brain, and simulated passenger cell fractions from a normal distribution. We considered two independent clones as well as a parent clone and a subclone (**Extended Data Figure 5a-d**). The simulations revealed that multiple clones with the same apparent size within the blood, such as those present in UW1, must represent independent clones, due to the lack of shared founding passengers. Secondly, these data highlighted that the infiltration of multiple independent clones will be underestimated by bulk data, due to the additive nature of the clones (**Extended Data Figure 5**). Finally, we considered three potential models for clonal competition (**Extended Data Figure 5e**), in which competing clones equally enter the brain (neutral model), the younger clone colonizes the brain more (early bird model), or the older clone evolves an enhanced ability to colonize the brain (late bloomer model). We find evidence for both neutral competition (e.g., ACT5 and ACT6) as well as the early bird model (e.g., UW3).

Finally, we searched for potential SNV or INDEL driver mutations not detected by whole genome sequencing by performing targeted sequencing of a known list of SNV or INDEL exons (**Supplementary Table 3**). We identified putative driver variants consistent with the primary PACT clone for four individuals as well as several CH-associated variants outside of the primary clone of other individuals that were not detected by blood WGS (**Extended Data Figure 6**; **Supplementary Table 4**). The brain specific mutations identified here may be marrow-derived driver variants with a low cell fraction in the blood or may represent variants independently evolved within the yolk sac-derived microglia^33,34^. These results indicate that diverse types of clones present in the bone marrow share the ability to enter the human brain during aging and provide a conceptual framework for interpreting passenger-assisted clone-tracking data.

### Phenotype of infiltrating cells

To identify the brain cell type harboring these HSC-derived PACT mutations, we performed single nucleus sequencing of UW1-UW16. We also included 3 individuals with a lower passenger burden, designated “low CH”, UW17-UW19. We obtained 141,525 nuclei from UW1-19 which we clustered together with an additional 44,550 nuclei from external reference brain and blood data^35,36^, for a total of 186,075 nuclei (**Figure 2a-c**). We identified 6 brain cell types, including microglia, and 3 blood cell types, including T cells, B cells, and monocytes (**Figure 2b**; **Extended Data Figure 7**). In all brain samples, including individuals with CH, low CH controls, and external reference samples, 99.35% of all hematopoietic brain nuclei clustered as *SALL1*+, *TMEM119*+ microglia (**Supplemental Table 5**). Examining the microglia cluster by sample, we found that all samples exhibited accessibility at the *SALL1* and *TMEM119* loci but not at the *ANPEP* locus, consistent with reference microglia and inconsistent with blood monocytes (**Figure 2d**). Comparing the median PACT cell fraction to the proportion of microglia present in each sample, we estimate that an average of 32.41% of the microglia consist of infiltrating cells with UW6 and UW16 having 85.14% and 92.67%, respectively (**Figure 2e**; **Supplemental Table 6**).

**Figure 2:**
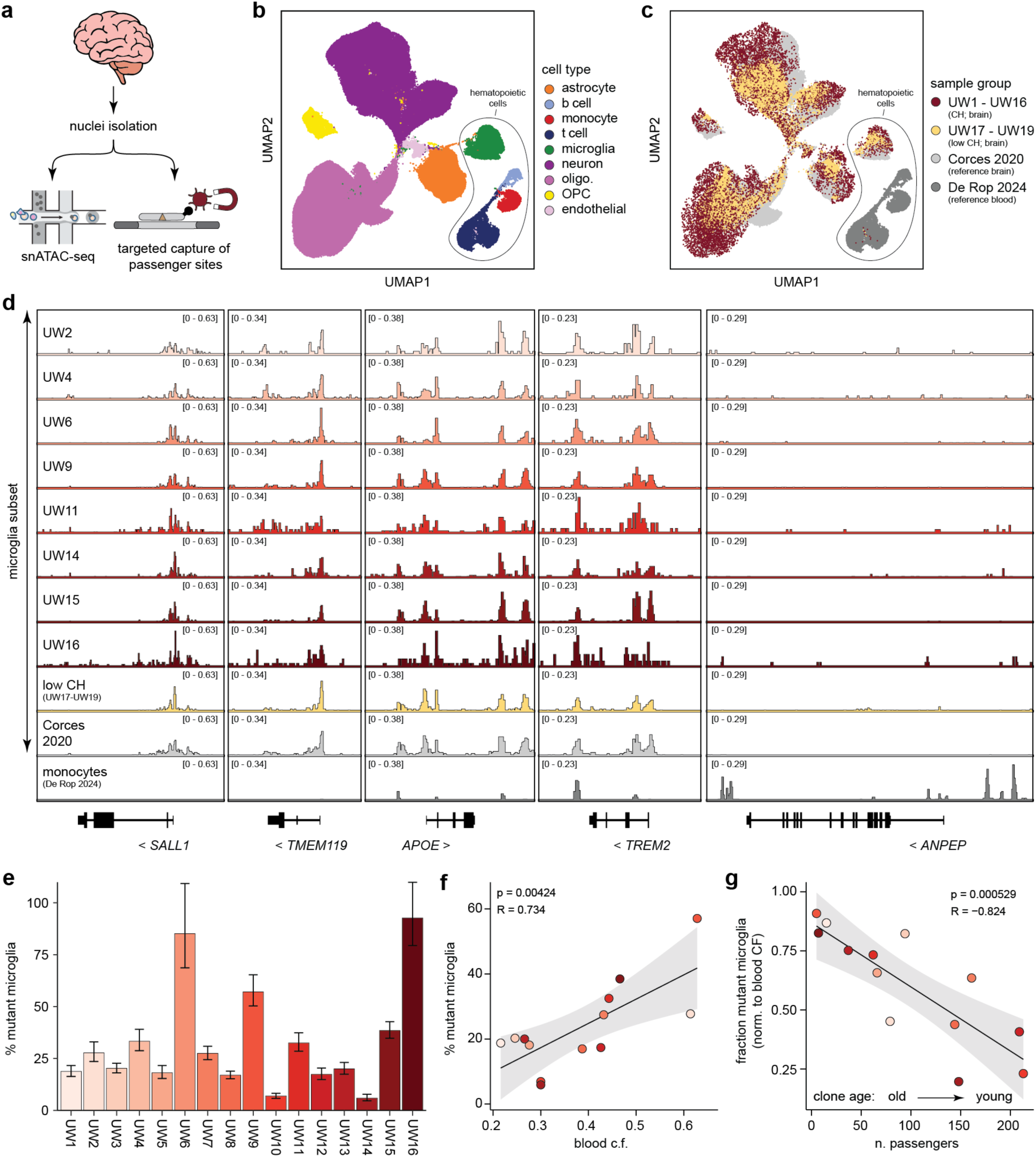
Single nucleus sequencing demonstrates that infiltrating cells are similar to microglia. **(a)** Schematic of experimental workflow. Single nuclei suspensions were prepared from postmortem brain specimens. Each sample was subjected to deep sequencing of selected variant sites and single nucleus sequencing in parallel. **(b)** snATAC-seq profiles of nuclei colored by cell type assignment. **(c)** snATAC-seq profiles of nuclei colored by sample group of origin. CH and low CH sample groups were down sampled to 10,000 nuclei per group for visualization. **(d)** Pseudobulk snATAC-seq tracks for selected gene loci. Reference microglia from Corces 2020 and reference blood monocytes from De Rop 2024 are also included. **(e)** Estimation of the proportion of microglia derived from the PACT clone in each individual. Error bars indicate the simulated 95% confidence interval for the estimate in each sample using n=10^6^ random samples. **(f)** Correlation between the percent mutant microglia and the cell fraction of the PACT clone in the blood. **(g)** Correlation between the fraction mutant microglia, normalized to blood clone size, and the number of passenger mutations present on the clone.

We re-clustered 7,889 myeloid cells, including the microglia from UW1-19 and the reference blood monocytes to investigate whether all myeloid cells within these brain samples were genuine microglia (**Extended Data Figure 8**). We identified 3 sub-clusters originating in the blood and 6 brain myeloid sub-clusters. All brain myeloid sub-clusters exhibited high accessibility of microglia hallmark genes *SALL1*, *NAV3*, and *TMEM119*, and no brain myeloid sub-cluster exhibited accessibility at markers of other brain macrophage subtypes, such as *CD163*, *MRC1*, or *HLA-DRA* (**Extended Data Figure 8e**). Secondly, we identified LoY and WT (chrY+) nuclei within our snATAC-seq data for the individuals with LoY clones present in their blood (UW4; UW12-UW16; **Extended Data Figure 9a-c**). The nuclei with LoY were predominantly assigned to the microglia cell type (average percent of nuclei with LoY = 26.2% for microglia versus 0.34% for other brain cells; **Extended Data Figure 9a**). The percentage of mutant microglia estimated via single nucleus LoY detection was correlated with our bulk estimates (R=0.650; **Extended Data Figure 9d**; **Supplementary Table 6**). Examining the microglia subclusters represented in each donor in nuclei with and without LoY revealed that the LoY and WT nuclei typically had similar cluster distributions (**Extended Data Figure 9e-f**). One exception was the myeloid sub-cluster C9, which appeared to consist of oligodendrocyte-microglia doublets and was exclusively represented by the WT nuclei (**Extended Data Figure 9e-f**). The lack of LoY-assigned nuclei in C9 may be due to technical factors, because a doublet containing a chrY+ oligodendrocyte would obscure the LoY status of the associated microglial nucleus. These results suggest that infiltrating cells adopt a phenotype very similar to most or all subtypes of endogenous microglia.

We sought to identify features predictive of microglia supplementation. For this analysis, we excluded the three individuals (UW4, UW6, and UW16) who had a larger fraction of mutant microglia than their blood cell fraction, reasoning that idiosyncratic factors may be involved in the disproportionate level of microglia supplementation observed in these three subjects. Considering the other 13 subjects, we found a positive correlation between clone size in the blood and percent mutant microglia (**Figure 2f**). Furthermore, quantifying clone age by counting the number of passengers on each primary clone, we found that older clones had a higher percentage of mutant microglia, after adjusting for the blood clone size (**Figure 2g**). These findings support the “early bird model” in which older clonal expansions have a competitive advantage in colonizing the brain due to a more prolonged seeding or longer residence time.

Finally, we sought to investigate how microglia supplementation might vary during human aging. For this analysis, we identified four additional individuals to perform PACT on, three in their 70s and one aged individual (UW20-UW23; **Supplementary Table 2**). We performed nuclei sorting to identify informative passenger mutations as well as single nuclei sequencing to identify cell phenotypes (**Extended Data Figure 10a-c**). We clustered 36,454 cells from UW20-23 with the low CH controls and external brain and blood reference datasets as described above, for a total of 93,307 nuclei, and estimated the percent mutant microglia in each sample (**Extended Data Figure 10b-e**; **Supplementary Table 7**). As before, we found that blood clone size was predictive of the percent mutant microglia, but with a lower extent of infiltration for each blood clone size – for example, UW22 had a cell fraction of 71.8% in the blood but only 22.6% mutant microglia (**Extended Data Figure 10f**). After controlling for blood clone size, individuals more than 80 years old had more mutant microglia than those younger than 80 years of age (**Extended Data Figure 10g**). When taken together with the fact that hematopoietic clone size also increases with age, these results suggest that microglia supplementation can begin in septuagenarians, if not earlier, and continues to increase during human aging, with infiltrating cells ultimately comprising a substantial fraction of the microglial pool in aged individuals.

### mtDNA lineage tracing of two research autopsies

To investigate the single-cell phenotype of the brain-infiltrating blood cells using an orthogonal approach, we sought to use lineage tracing based on mitochondrial DNA (mtDNA) variants^28,29,37,38^. Unlike nuclear mutations, the high copy number of mtDNA enables robust detection of mutations in individual cells^28,29,37,38^. We obtained fresh tissue specimens from two research autopsies and performed mitochondrial single-cell assay for transposase-accessible chromatin with sequencing (mtscATAC-seq; **Figure 3a**)^38^. The first autopsy specimen, RA1, was a 61-year-old female who was diagnosed with a myeloid neoplasm and underwent a bone marrow transplant (BMT) 22 months before passing away. The second autopsy specimen, RA2, was a 77-year-old male with coronary artery disease whose cause of death was a right middle cerebral artery stroke. From both specimens, we obtained fresh brain tissue, digested the tissue to obtain intact cells, performed flow cytometry to isolate high-quality live cells, and then performed mtscATAC-seq. From RA2, we also performed mtscATAC-seq on cells isolated from the bone marrow.

**Figure 3:**
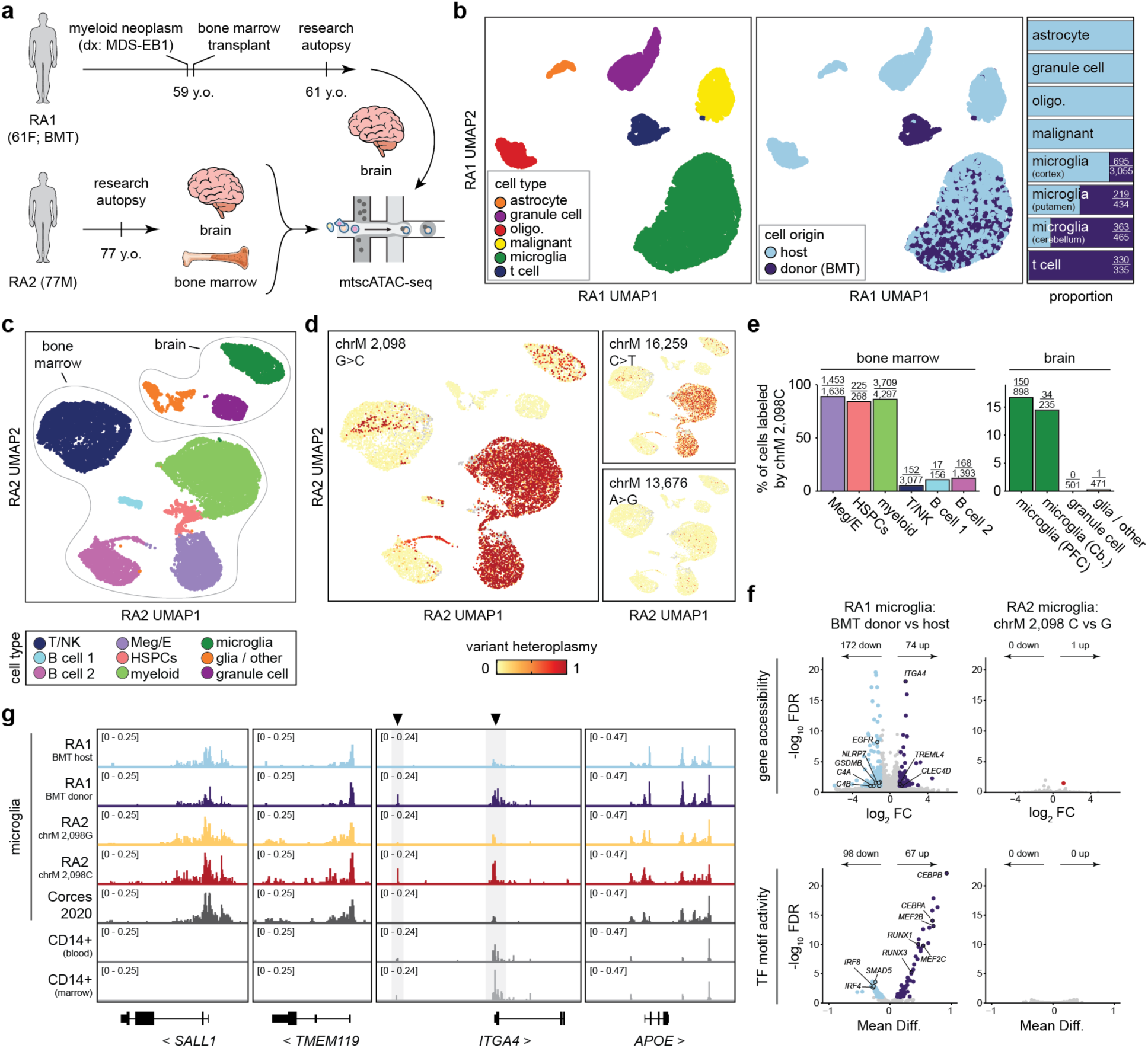
Infiltrating marrow-derived cells are almost identical to microglia in a 77-year-old male. **(a)** Schematic of experimental workflow. **(b)** scATAC-seq profiles of cells isolated from three brain regions of RA1 colored by cell type assignment (left) or by genotype (center). Quantification of the proportion of each genotype within each cell type is shown on the right and is additionally broken down by brain region for microglia. **(c)** scATAC-seq profiles of cells isolated from two brain regions and the bone marrow of RA2 colored by cell type assignment. **(d)** scATAC-seq profiles of RA2 cells colored by mtDNA variant heteroplasmy in each cell. **(e)** Quantification of the number of RA2 cells labeled by variant chrM 2,098C in each cell type in the marrow (left) and brain (right). **(f)** Differential analysis comparing genotypes within the microglia cluster of RA1 (left) or RA2 (right). Differentially accessibly genes (|Log_2_FC| > 1 and FDR < 0.1) and differentially active TF motifs (|Mean Diff.| > 0.01 and FDR < 0.1) are both shown. **(g)** Pseudobulk scATAC-seq tracks for selected gene loci. BMT, bone marrow transplant; HSPCs, hematopoietic stem and progenitor cells; Meg/E, megakaryocyte/erythroid; Cb, cerebellum; PFC, pre-frontal cortex.

For RA1, we sequenced 6,239 cells from 3 brain regions, including the cortex, cerebellum, and putamen, and identified 6 distinct cell types (**Figure 3b**; **Extended Data Figure 11a-c**). Donor and host cells could be easily distinguished by using genetic demultiplexing based on nuclear germline polymorphisms^39,40^, which were also concordant with mitochondrial germline polymorphisms detectable in the mtscATAC-seq data (**Figure 3b**; **Extended Data Figure 11d-e**). As expected, the non-hematopoietic clusters exclusively consisted of host cells. Within the hematopoietic cells, we identified a cluster of T cells, which was derived from the donor, and microglia, which had been partially replaced by donor cells (**Figure 3b**). We next investigated 14,227 cells from the bone marrow and brain of RA2, including both the pre-frontal cortex and the cerebellum (**Extended Data Figure 12a-b**). We identified 6 main cell types in the bone marrow and 3 in the brain, including microglia (**Figure 3c**; **Extended Data Figure 12c**). We examined the mtDNA mutations in this individual, seeking to identify mutations that could specifically label bone marrow derived cells (**Extended Data Figure 12d-e**). We identified three mutations matching these criteria, which almost exclusively labeled the myeloid lineage of the bone marrow – including hematopoietic stem and progenitor cells (HPSCs), megakaryocytes / erythrocytes, and CD14+ cells in the marrow – and were also present in the microglia cluster in the brain but were less abundant in other immune lineages and essentially absent in the non-hematopoietic brain cell types (**Figure 3d**). Strikingly, one of these mutations, chrM 2,098G>C, labeled 83.96% of the HSPCs and 86.32% of the myeloid cells in the bone marrow, serving as an ideal marker for marrow-derived myeloid cells. ChrM 2,098C labeled 14.47% and 16.70% of the microglia in the cerebellum and the pre-frontal cortex of RA2, respectively (**Figure 3e**).

We sought to identify differences between the infiltrating and endogenous microglia in the two subjects. Unbiased analysis of chromatin accessibility, gene accessibility and transcription factor motifs identified modest differences within the microglia derived from the BMT in RA1 (**Figure 3f**; all significant differential items are reported in **Supplementary Tables 8-11**). In RA2, one gene reached genome-wide significance (**Figure 3f**; **Supplementary Table 11**). *ITGA4* was the top gene marking donor derived microglia in RA1^41,42^. Next, we visualized chromatin accessibility at several marker genes associated with microglial identity for both microglial genotypes of RA1 and RA2, previously published reference human microglia, and CD14+ myeloid cells from the blood and bone marrow (**Figure 3g**). In RA1, both genotypes appeared indistinguishable at important microglia gene loci *SALL1*, *TMEM119*, and *APOE*. Given that this donor had BMT only 22 months prior, this suggests that engrafting cells can adopt a microglial phenotype in the human brain in a relatively short time frame. Increased accessibility at two regions within the *ITGA4* locus were apparent in infiltrating cells in both RA1 and RA2, that were less accessible in the other microglia subsets. We also analyzed other key genes, including previously reported ontogeny markers and genes defining specific brain microglia or macrophage subtypes, and did not find additional consistent differences between the infiltrating myeloid cells and microglia (**Extended Data Figure 13**)^22,24,43^. These results indicate that infiltrating cells adopt nearly identical phenotypes to microglia, although small differences may remain, providing phenotypic clues which indicate the origins of these cells.

### Clonal hematopoiesis and Alzheimer’s disease

Our results indicated that the presence of peripheral myeloid cells in the brain parenchyma occurs in individuals with all types of CH. We wondered if other forms of CH might also be associated with protection from AD, beyond the protective association we previously reported between SNV or INDEL-driven CH and AD^5,16,17^. For these analyses, we used WGS data available from 20,379 participants from the ADSP with known age at blood draw used for WGS or known age at death if brain autopsy was performed. Each participant had a clinical AD dementia diagnosis based on criteria of the National Institute of Neurological and Communicative Disorders and Stroke (NINCDS) and the AD and Related Disorders Association (ADRDA) for definite, probable or possible AD, and a subset had the diagnosis confirmed by neuropathological assessment. For each subject, we identified SNV or INDEL mutations, mCAs, and passenger mutations. Since bona fide passenger mutations are only detectable in low-depth WGS if there is a clonal expansion event, we used a threshold count for passenger mutation burden to classify individuals into those with CH with unknown drivers versus no CH or other forms of CH (**Supplementary Table 12**). We found that all types of driver events increased with age (**Figure 4a**). In individuals over 90, 38.75% had SNV or INDEL drivers, 10.5% had mCA drivers, and 28.0% had an unknown driver. In aggregate, the prevalence of all types of CH increased from 10.2% in individuals under 60 (mean age: 54.97 years old) to 77.25% in individuals at least 90 years old.

**Figure 4:**
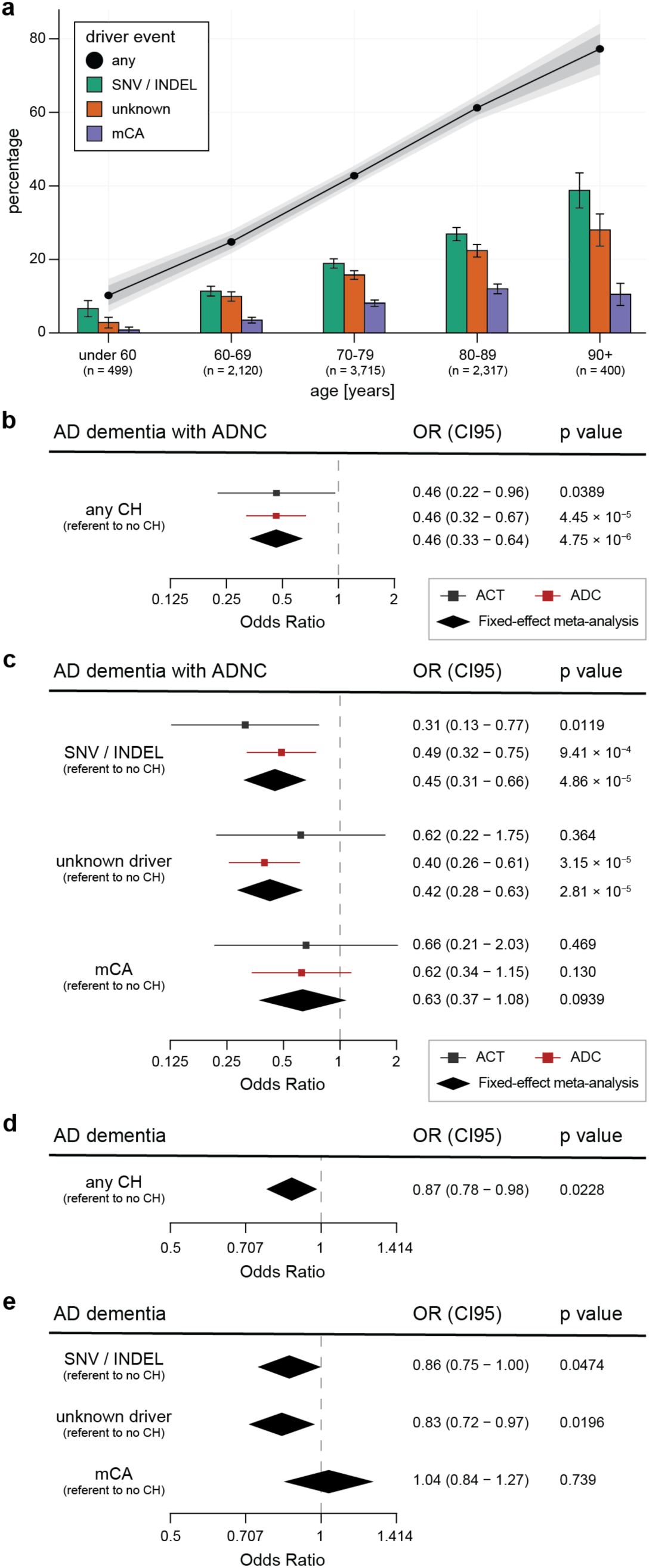
Clonal hematopoiesis is associated with reduced AD dementia and ADNC. **(a)** Prevalence of each type of somatic clonal expansion in blood during aging. Error bars indicate CI95. 95% and 99.9% confidence intervals are shown for the proportion of individuals with any driver event. n=9,071 subjects in the ADC cohort with WGS data and age at blood draw available. **(b)** Effect of CH on the risk of AD dementia with ADNC. **(c)** Effect of CH driver type on the risk of AD dementia with ADNC. **(d)** Fixed-effect meta-analysis for risk of AD dementia in CH carriers using logistic regression in six ADSP cohorts with two-sided Wald P value shown. **(e)** Fixed-effect meta-analysis for risk of AD dementia by driver type in six ADSP cohorts with two-sided Wald P value shown. **(b-c)** include n=192 subjects in the ACT cohort and n=1,273 subjects in the ADC cohort. Additional demographic information is available in Supplementary Table 3. OR, CI95, and two-sided Wald P value were calculated from logistic regression models which included age, sex, and *APOE* genotype as covariates. The measure of center is the OR and the lengths of the lines represent the CI95 for the OR. **(d-e)** include n=8,352 individuals from six ADSP cohorts. Additional demographic information is available in Supplementary Table 5. For meta-analyses, the center of the diamonds represents the OR and the lengths represent the CI95 for the OR. SNV/INDEL, single nucleotide variant or insertion or deletion; mCA, mosaic chromosomal alteration; OR, odds ratio; CI95, 95% confidence interval; AD, Alzheimer’s disease; ADNC, AD neuropathologic change; CH, clonal hematopoiesis.

We initially focused on the highest confidence AD cases and controls who have diagnoses confirmed by postmortem neuropathology (**Supplementary Table 13**). These represent “gold standard” AD diagnoses with excellent statistical power for association tests^44^. We used subjects within two ADSP cohorts with AD Neuropathologic Change (ADNC) data available, ACT and ADC (n=171 and n=1,133 neuropathologically confirmed diagnoses, respectively). We used a previously described composite ADNC score that quantifies the accumulation of the two neuropathological hallmarks of AD – β-amyloid plaques and tau neurofibrillary tangles – in combination with clinical dementia. We found that CH was associated with protection from neuropathologically confirmed AD using logistic regression (**Figure 4b**; all driver types aggregated; OR=0.45, p=1.8×10^−6^). Breaking down the individual types of driver events, we found that SNV or INDEL drivers and unknown drivers were individually associated with protection from AD, while mCA drivers were not significant (**Figure 4c**). Finally, we performed exploratory analyses on additional subsets of CH. In line with our prior results, *DNMT3A*, *TET2*, and other mutated genes showed similar effect sizes (**Supplementary Table 14**; **Extended Data Figure 14a**). We also found that for individuals with at least one SNV or INDEL driver, individuals with multiple driver events appeared to have the largest risk reduction (**Supplementary Table 14**; **Extended Data Figure 14b**).

To replicate this association in additional cohorts, we selected six ADSP cohorts with available information on NINCDS-ADRDA diagnoses of clinical AD dementia (**Supplementary Table 15**), including three case-control cohorts (ACT, ADC, and MIA) as well as three longitudinal cohorts (ASPREE, WHICAP, and ADNI). Again, we identified CH as being associated with reduced risk of AD (**Figure 4d**; **Extended Data Figure 15**). Both SNV or INDEL drivers and unknown drivers exhibited a protective odds ratio while there was no conclusive association between mCAs and AD (**Figure 4e**; **Extended Data Figure 15**). Overall, these results suggest a protective role for CH in AD progression, beyond individuals with known SNV or INDEL drivers.

## Discussion

Our data provide evidence that expanded clones of HSC-derived myeloid cells colonize the brain and supplement the microglial pool during homeostatic human aging. Our findings that infiltrating myeloid cells can adopt a phenotype almost indistinguishable to microglia are unexpected in light of prior studies, which required engineered or pathological settings to recruit peripheral myeloid cells into the brain and then found phenotypic differences between the infiltrating cells and resident microglia^22–24,45–49^. Examining individuals spanning three decades of the human lifespan, we find evidence that marrow-derived cells can be detected in the brain parenchyma as early as age 73 and become more prevalent in older individuals. By analyzing donor cells in a bone marrow transplant recipient, we find that the engrafted cells can adopt a microglial phenotype, including accessibility at the *SALL1* locus, within 22 months of residence. Our findings suggest that, in humans, residence in the brain enables monocyte-derived cells to adopt the full microglial phenotype, although minor differences may persist.

Because these invading cells adopt a phenotype nearly identical to endogenous microglia, proving their origins required the development of PACT, a lineage tracing method that uses the somatic nuclear passenger mutations that accumulate during aging to track expanded clones of hematopoietic cells as they migrate from the blood into the brain. Concomitant application of PACT and single-nucleus sequencing demonstrated that peripherally derived cells can comprise up to 92.7% of the microglial pool and revealed key features of clonal competition within the brain, demonstrating that larger clones and older clones tended to contribute more to the microglial pool. Detection of loss of the Y chromosome in individual nuclei suggests that infiltrating cells can supplement most or all microglia subtypes. Importantly, PACT is compatible with archival human tissue specimens, providing a tractable approach to study somatic mosaicism in healthy human tissues.

The discovery of cellular crosstalk between the blood and the central nervous system that becomes common (perhaps ubiquitous) in healthy human aging suggests therapeutic opportunities for neurological disorders. Microglia replacement has shown promise in pre-clinical models and recently, in humans, either to replace defective microglia^50–56^ or to deliver therapeutic molecules into the brain^57^, but the requirement to deplete existing microglia requires complex and/or invasive regiments. The appreciation of a natural process in which blood cells can infiltrate the human brain and supplement the microglial pool, without prior conditioning or depletion of the existing microglia, suggests microglia reinforcement as an alternate strategy for central nervous system delivery and therapeutics. Our findings of a protective association between clonal hematopoiesis and Alzheimer’s disease provides preliminary evidence that augmenting the microglial pool might help to pre-empt the onset of neurodegenerative disease in humans^5,16^. Strategies that modulate the ability of peripheral myeloid cells to infiltrate the brain may lead to durable and pre-emptive treatments for human neurological disorders.

## Extended Data Figure Legends

**Extended Data Figure 1:**
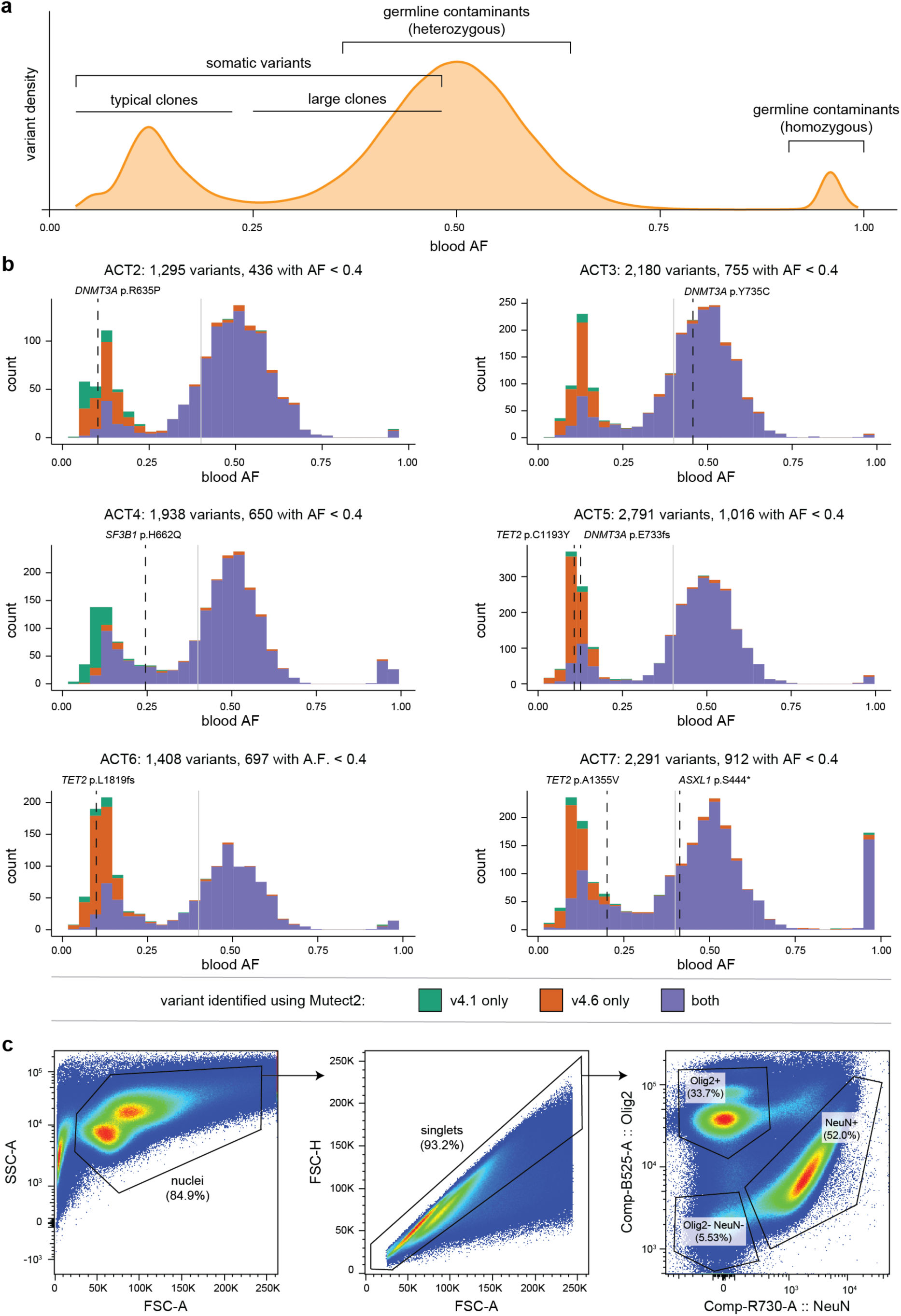
Variant calling and sort scheme for PACT. **(a)** Schematic of typical variant calling results in human blood by allele frequency. **(b)** Variant calling results for each ACT subject by allele frequency (AF). Cutoff for inclusion in the PACT panel (AF < 0.4) is shown in gray. Known SNV/INDEL driver(s) are shown as dashed lines. **(c)** Representative sort scheme for sorting human brain nuclei for PACT.

**Extended Data Figure 2:**
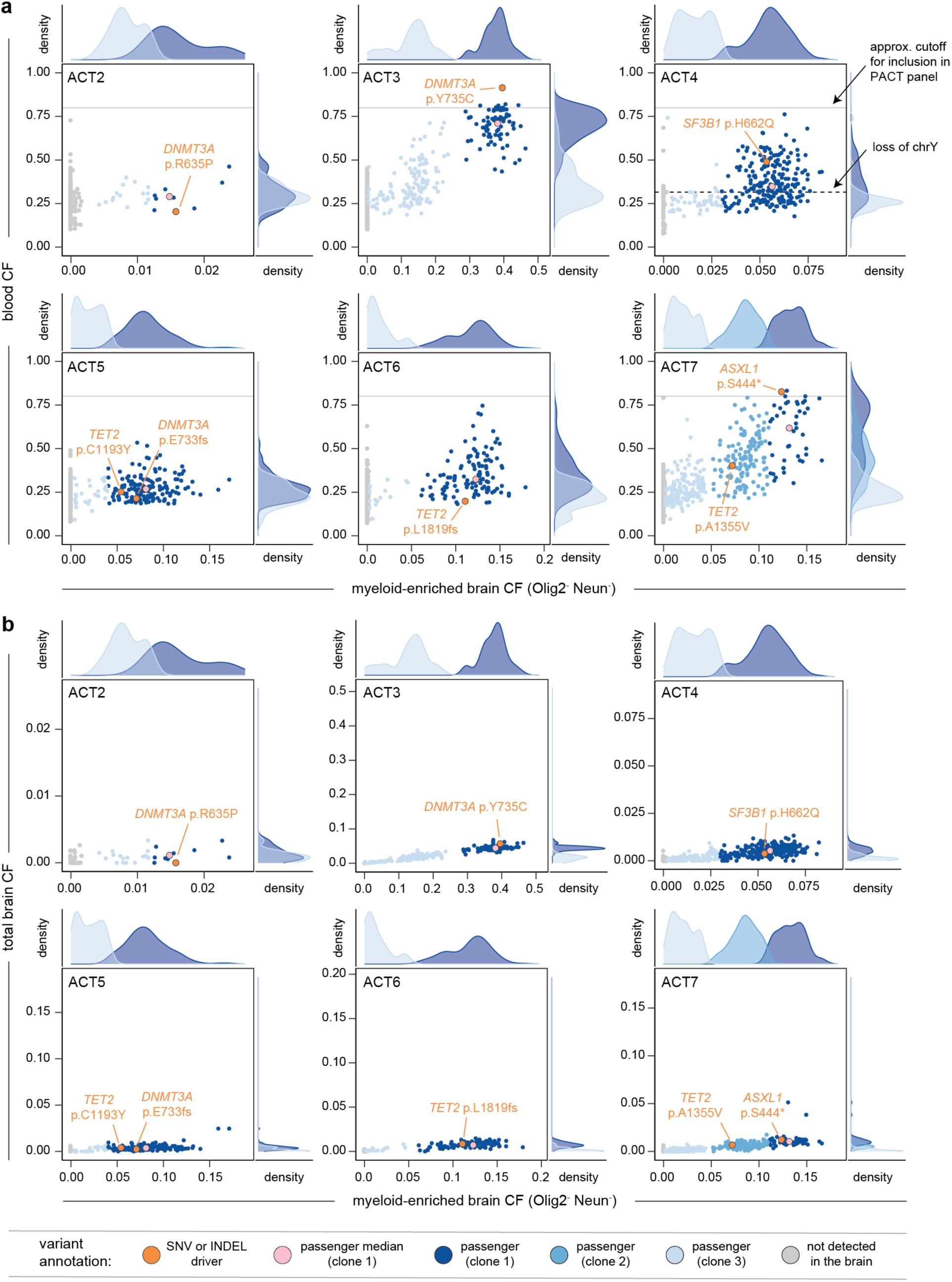
Validation of PACT using brain specimens from individuals with SNV or INDEL drivers. For each sample ACT2 – ACT7, the cell fraction of each passenger mutation in the myeloid-enriched fraction of the brain is plotted against the **(a)** blood cell fraction or **(b)** total brain cell fraction. The median cell fraction for the primary clone of passenger variants is shown in pink and known driver variants are shown in orange.

**Extended Data Figure 3:**
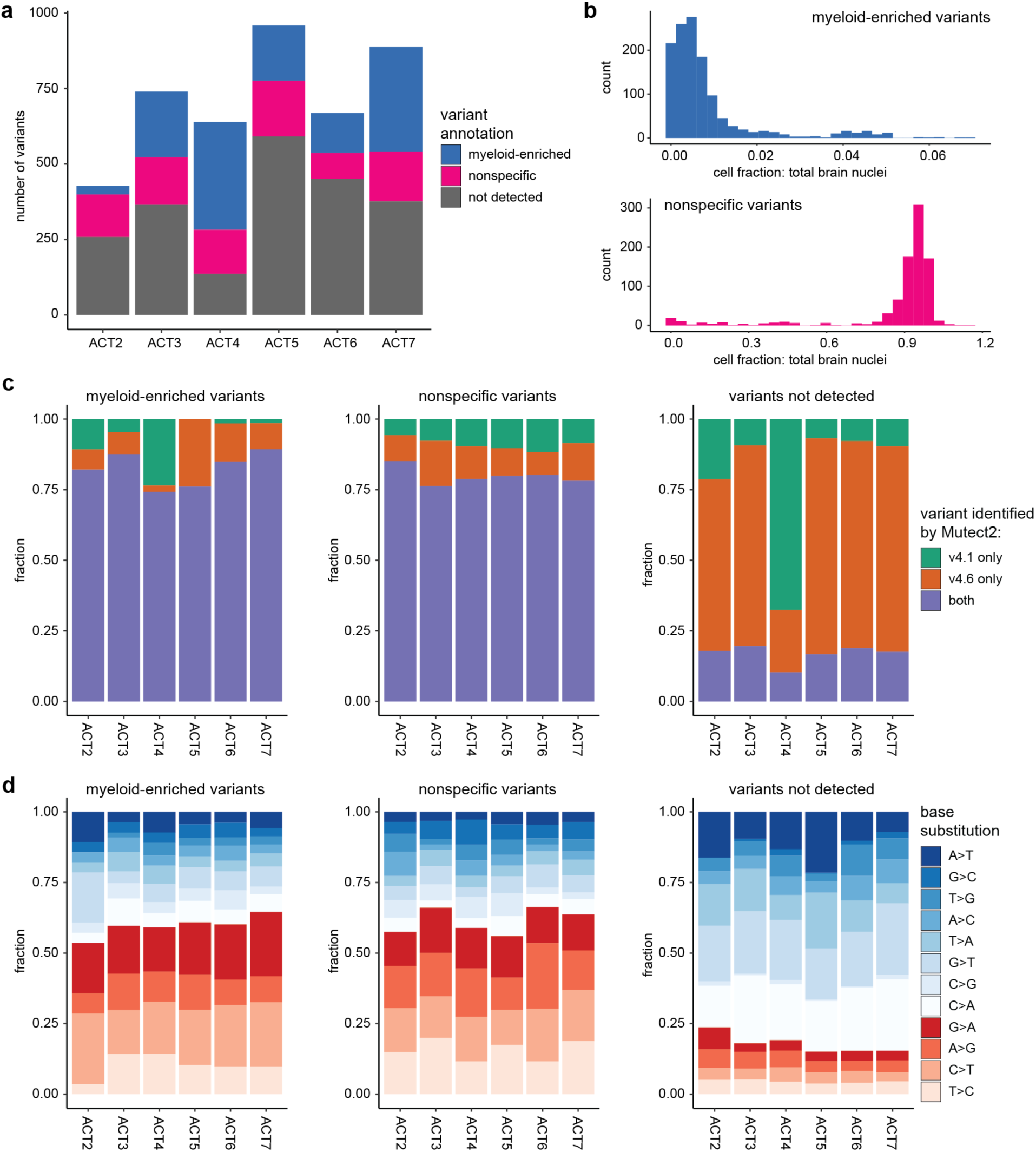
Retrospective analysis of informative variants for PACT. **(a)** Number of mutations identified as myeloid-enriched, nonspecific, or not detected from PACT results of each individual. **(b)** Histogram of the brain cell fractions of PACT variants for myeloid-enriched variants (top) and nonspecific variants (bottom). **(c)** Analysis of Mutect2 performance for each variant category and each individual. **(d)** Analysis of base substitutions (transitions or transversions) for each variant category and each individual.

**Extended Data Figure 4:**
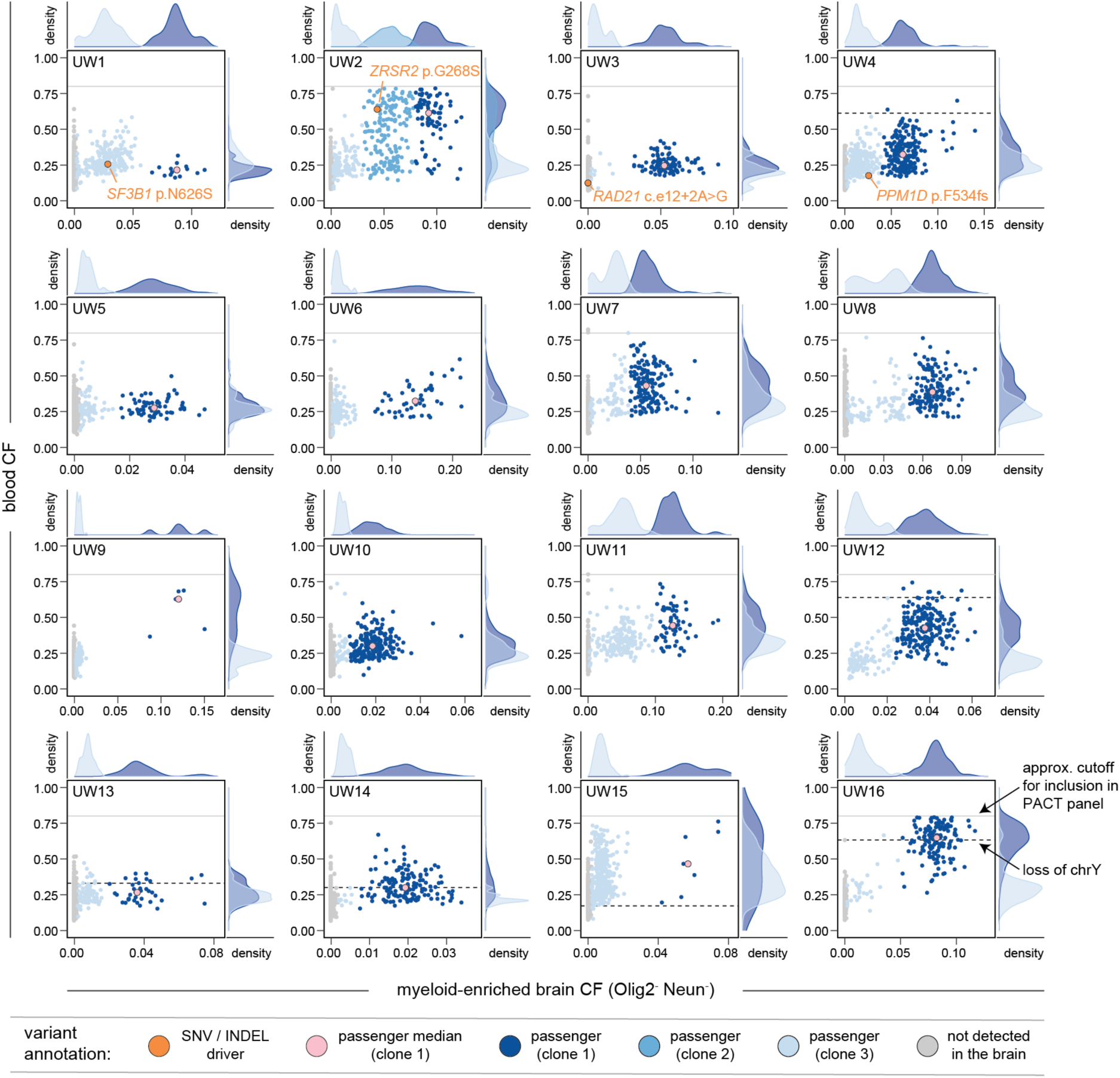
Empirical passenger distributions for marrow-derived clones infiltrating the human brain. For each sample UW1 – UW16, the cell fraction of each variant in the myeloid-enriched fraction of the brain and the blood is plotted. The median cell fraction for the primary clone of passenger variants is shown in pink and known driver variants are shown in orange.

**Extended Data Figure 5:**
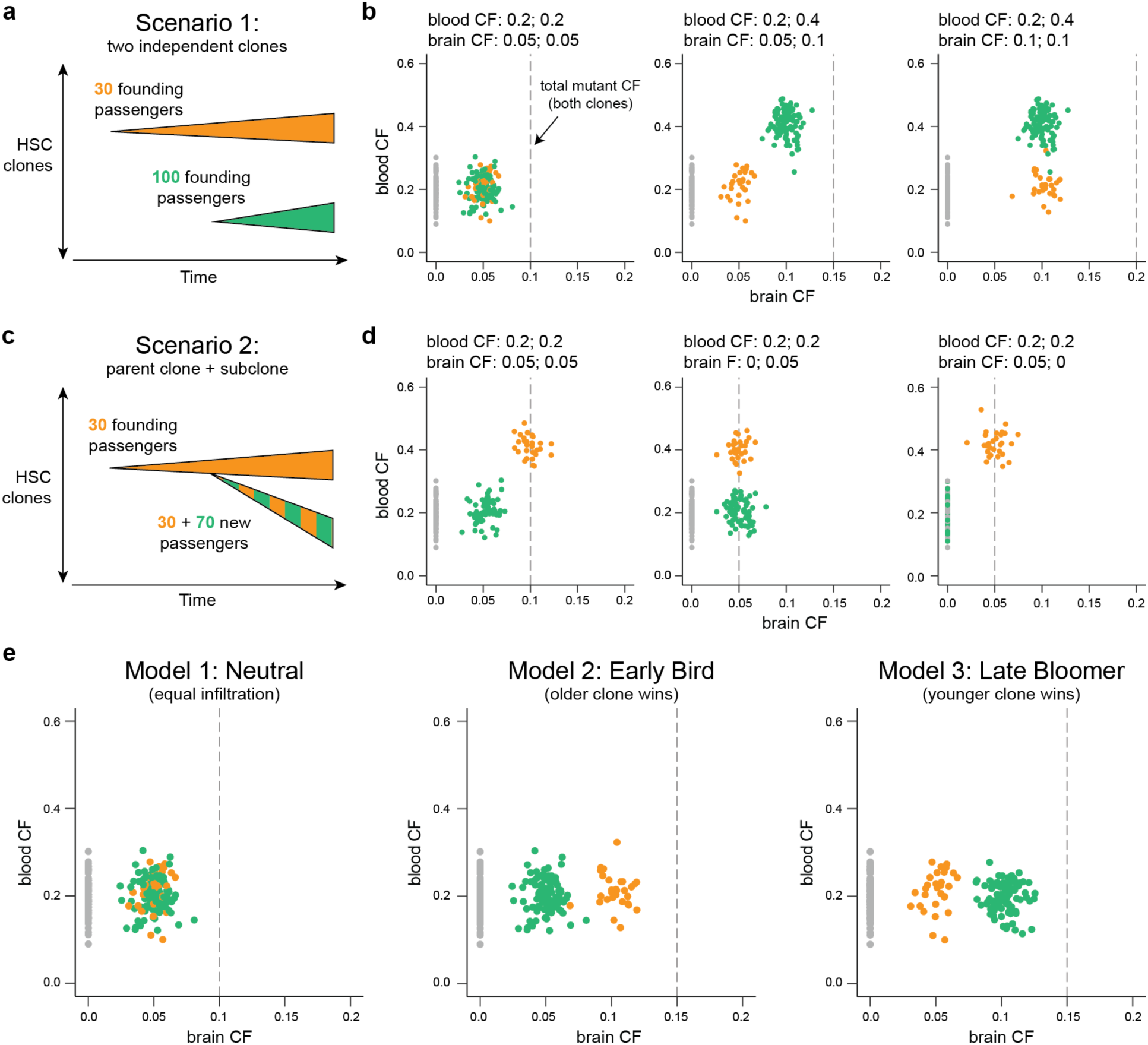
Simulated passenger distributions for different clonal evolution scenarios. **(a)** Illustration of two independent clones. **(b)** Simulated passenger distributions for two independent clones in a neutral scenario (left, middle) or when one clone is disproportionally represented within the brain (right). **(c)** Illustration of parent clone and subclone. **(d)** Simulated passenger distributions for a parent clone and subclone where both clones (left), subclone only (middle), or parent only (right) enter the brain. **(e)** Three models for clonal competition within the brain between independent HSC-derived clones that are equally expanded within the blood. **(b,d,e)** Gray dashed line indicates the total infiltrating cell fraction in the brain aggregated across both clones.

**Extended Data Figure 6:**
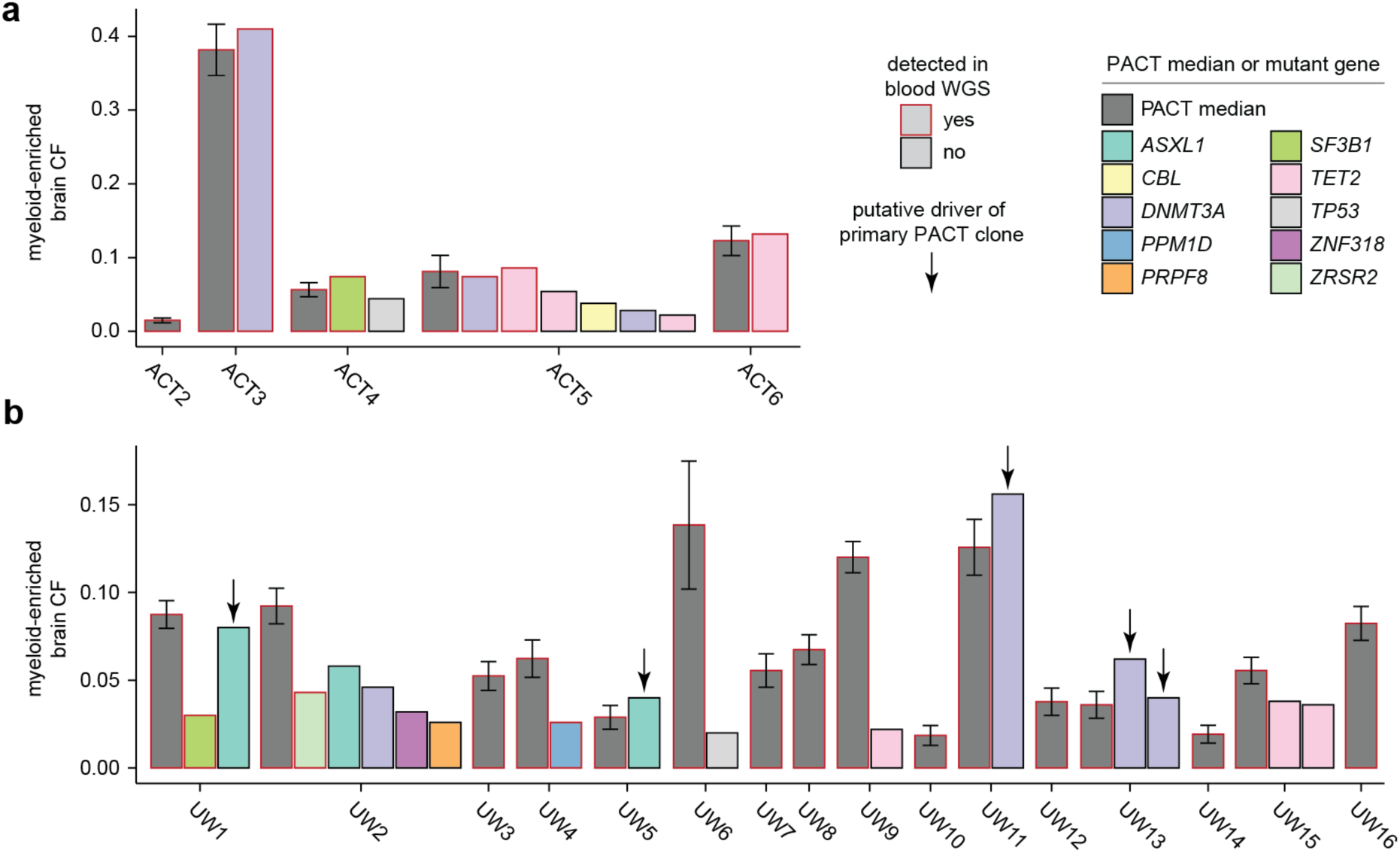
Variant calls from myeloid-enriched brain samples using a targeted panel of known CH drivers. The CF of individual variants are shown for **(a)** ACT2-6 and **(b)** UW1-16. Variants were manually curated and filtered based on a cell fraction of at least 0.02. The PACT median CF is included for reference. Error bars represent median ± median absolute deviation.

**Extended Data Figure 7:**
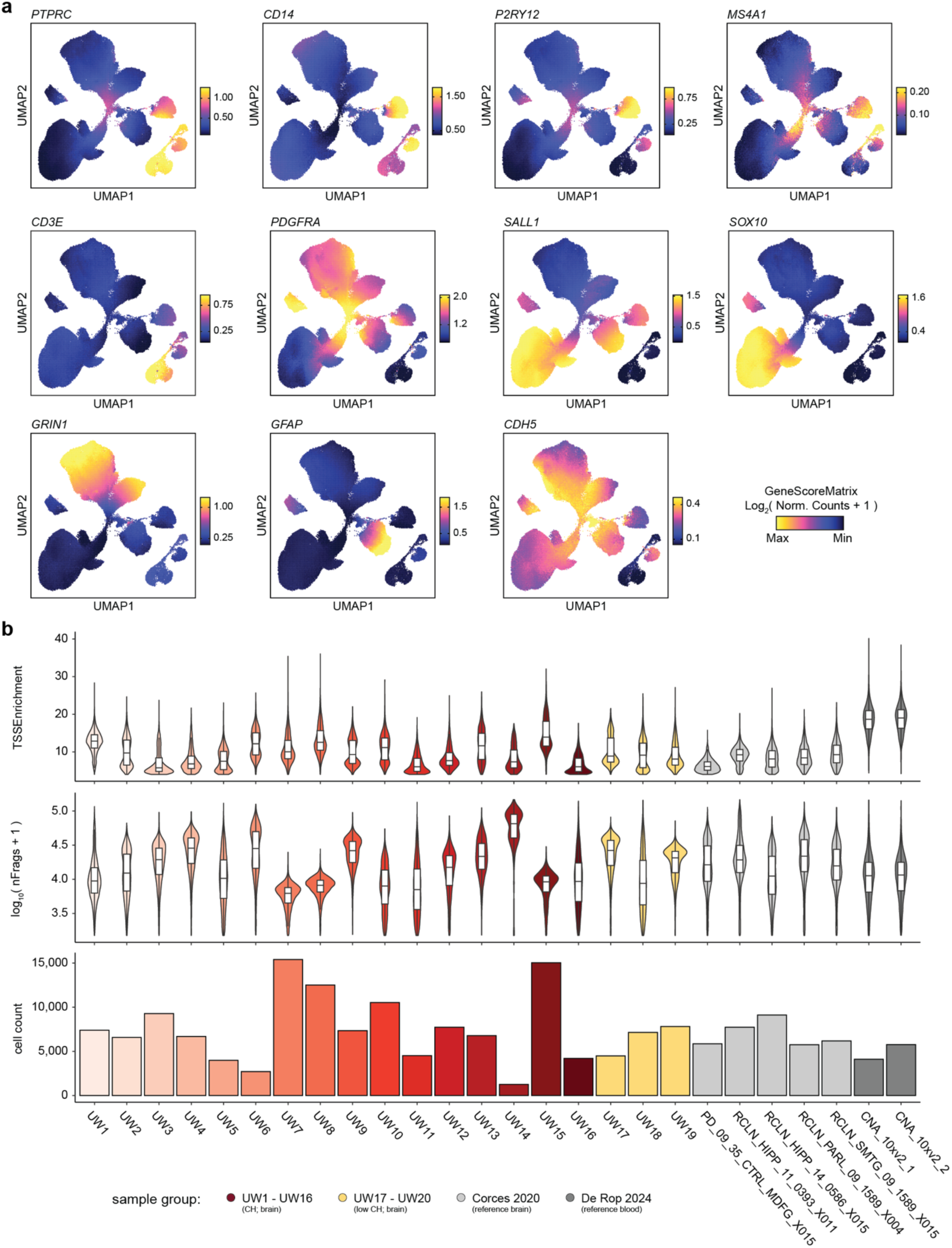
Marker genes and quality control for snATAC-seq data. **(a)** snATAC-seq profiles of nuclei colored by gene accessibility for selected marker genes. Gene accessibility is quantified using ArchR GeneScore. **(b)** snATAC-seq metrics for each sample in the present study, as well as each sample included from brain and blood reference datasets. TSS enrichment per cell (top), number of fragments per cell (center), and number of cells (bottom) are shown for each sample.

**Extended Data Figure 8:**
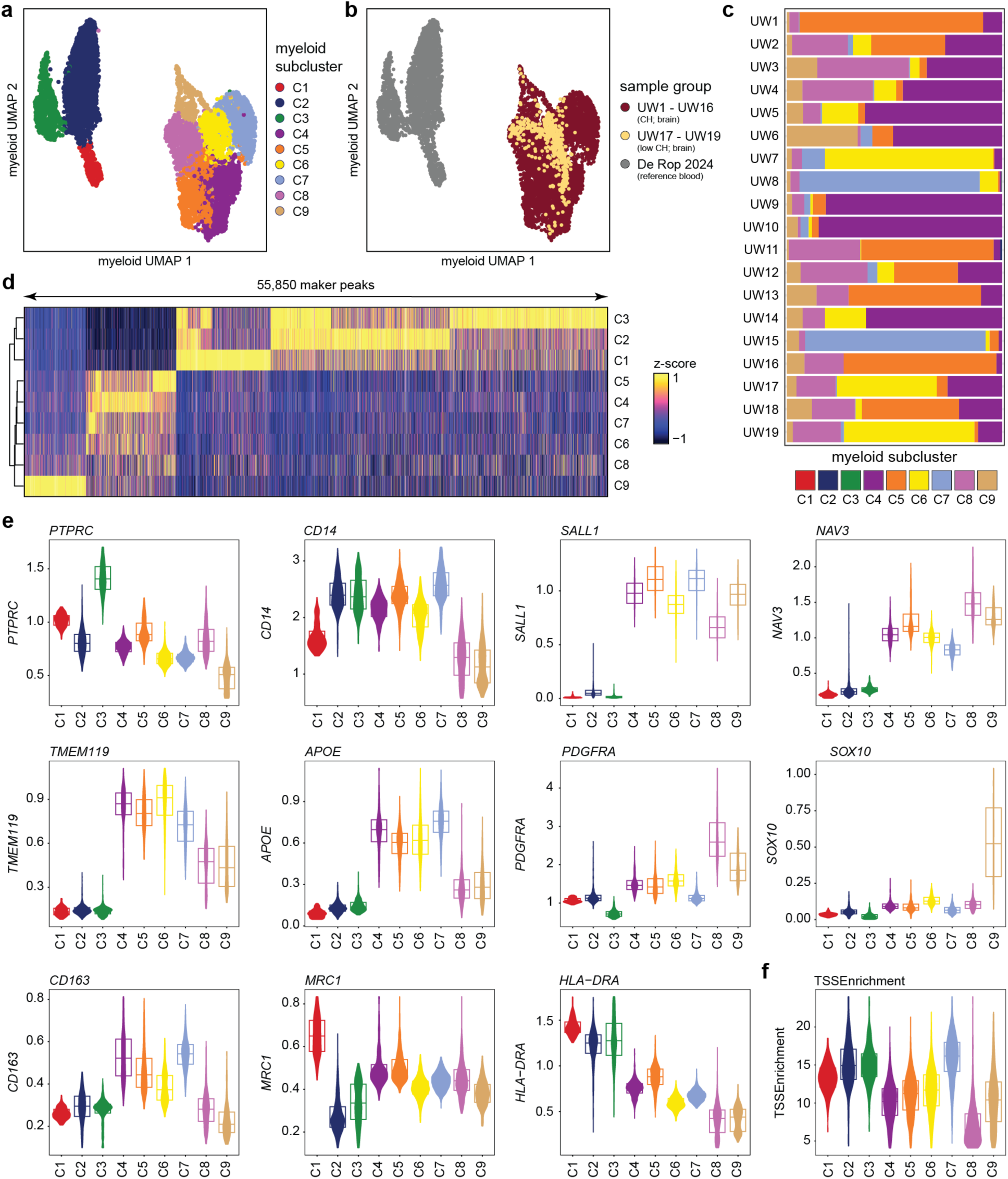
Sub-clustering of brain and blood myeloid cells. **(a)** snATAC-seq profiles of nuclei colored by myeloid sub-cluster. **(b)** snATAC-seq profiles of nuclei colored by sample group. **(c)** Quantification of the myeloid sub-cluster composition of each brain sample UW1-19. **(d)** Heatmap of marker peaks for each sub-cluster. Marker peaks were identified as FDR ≤ 0.05 and log_2_ (fold change) ≥ 1 in ArchR using the Wilcoxon test to compare cells in each sub-cluster. **(e)** Quantification of gene accessibility in each myeloid sub-cluster for selected marker genes. Gene accessibility is defined using ArchR GeneScore. **(f)** TSS enrichment of each cell in each myeloid sub-cluster.

**Extended Data Figure 9:**
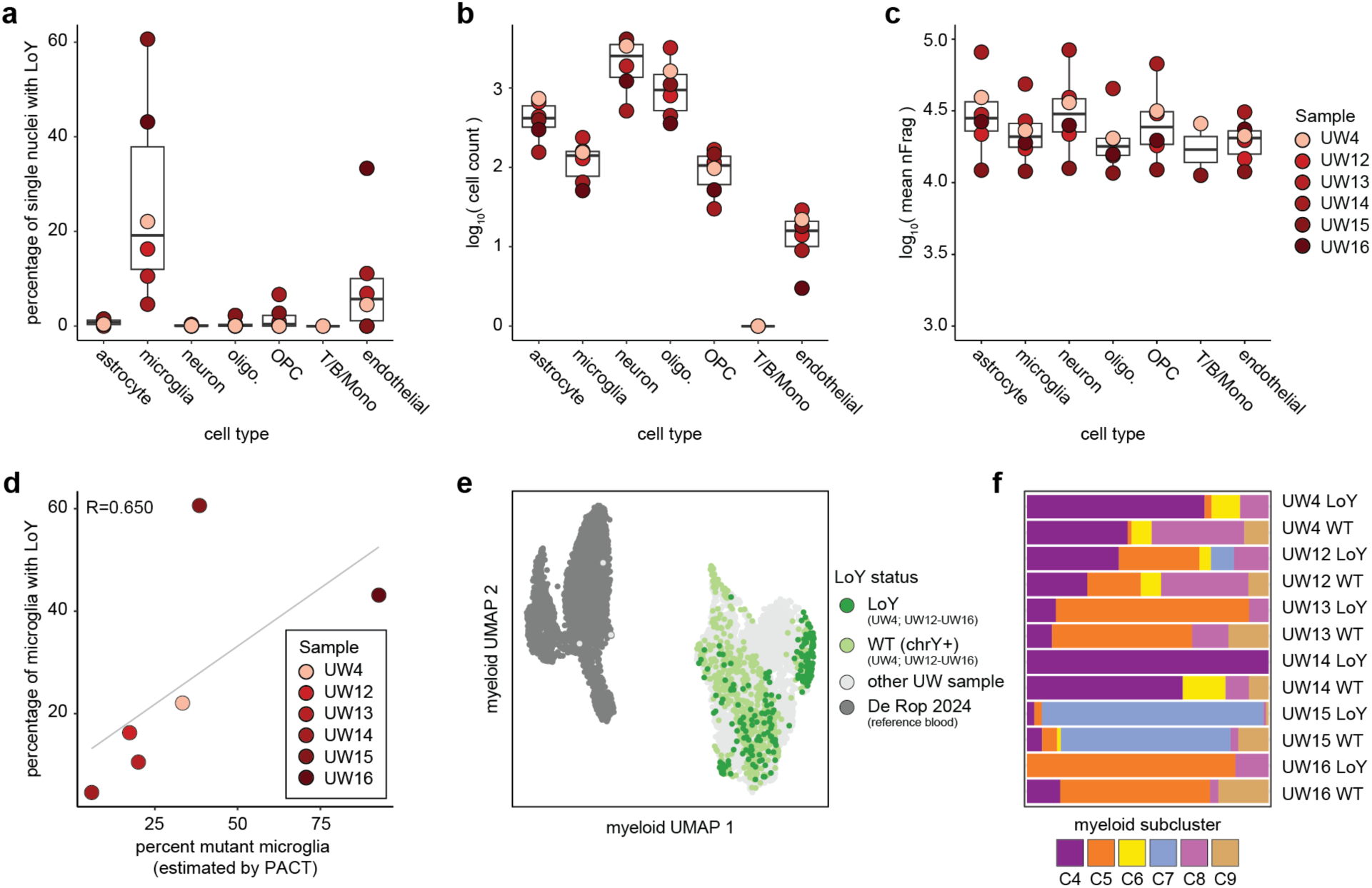
Detection of LoY in single nuclei from snATAC-seq data. **(a)** Percent of LoY single nuclei by cell type and sample. **(b)** Number of cells included in the LoY analysis for each cell type and sample. **(c)** Mean fragment count for each cell type and sample. Correlation between the percent mutant microglia estimated by PACT and the percent of microglia with LoY in single cells. **(e)** Individual LoY and WT nuclei visualized on the myeloid sub-clustering dimensionality reduction. **(f)** Quantification of the myeloid sub-cluster composition for LoY and WT nuclei within each sample. **(a-f)** LoY calling was performed in n=26,435 nuclei from samples UW4 and UW12-16 that had adequate sequencing depth (nFrags ≥ 10,000).

**Extended Data Figure 10:**
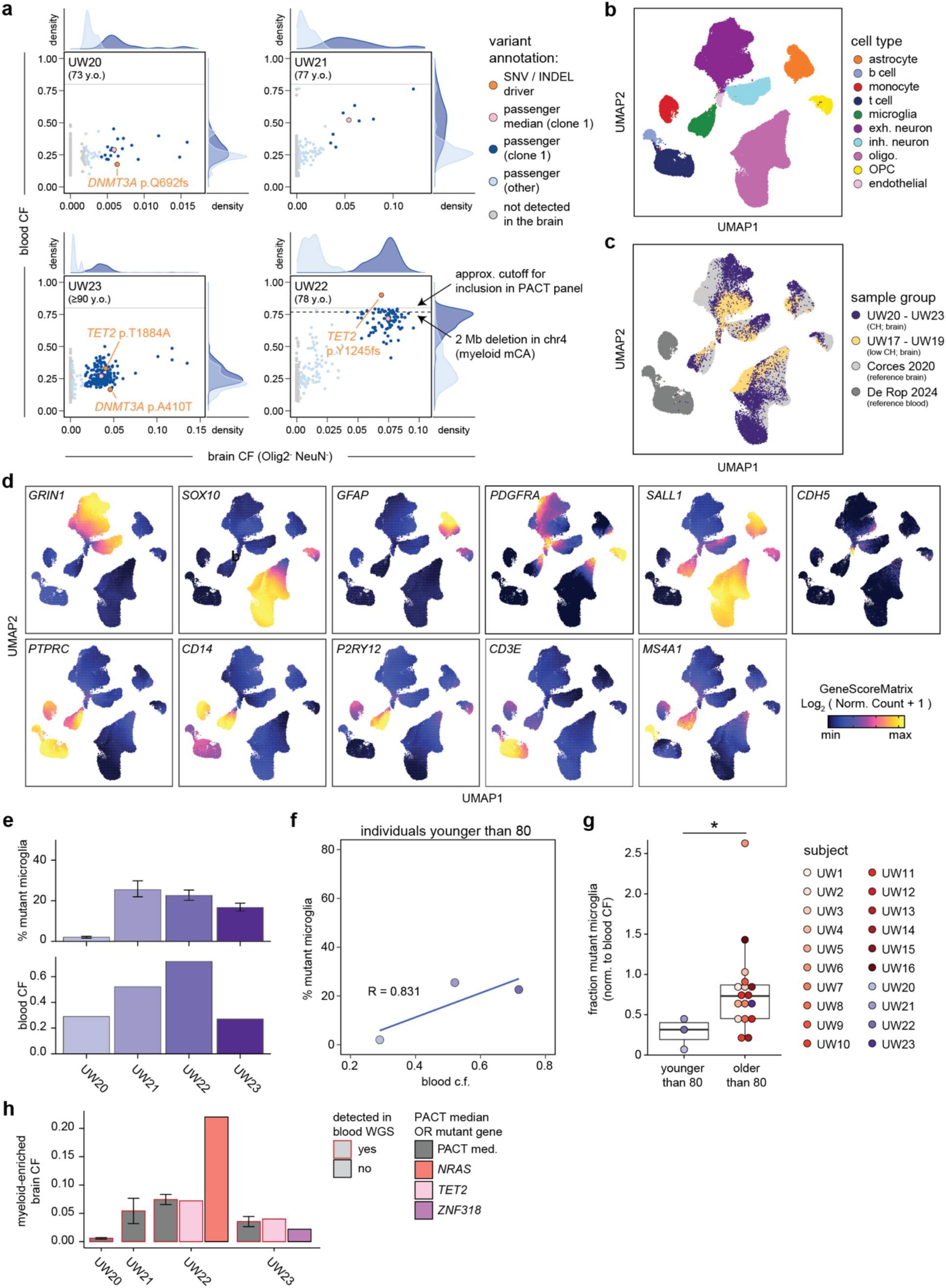
Microglia supplementation from the bone marrow increases with age. **(a)** Empirical passenger distributions for individuals UW20 – UW23. **(b)** snATAC-seq profiles of nuclei colored by cluster assignments. **(c)** snATAC-seq profiles of nuclei colored by sample group. **(d)** snATAC-seq profiles of nuclei colored by gene accessibility for selected marker genes. **(e)** Percent mutant microglia (top) and blood cell fraction (bottom) for individuals UW20 – UW23. Error bars indicate the simulated 95% confidence interval for the percent mutant microglia estimate in each sample using n=10^6^ random samples. **(f)** Correlation between percent mutant microglia and blood cell fraction for individuals younger than 80 (UW20 – UW22). **(g)** Comparison of percent mutant microglia (normalized to blood cell fraction) in individuals younger or older than 80. One-sided Wilcoxon rank sum exact test p=0.027 with n=20 donors. **(h)** Variant calls from myeloid-enriched brain samples using a targeted panel of known CH drivers. Variants were manually curated and filtered based on a cell fraction of at least 0.02. The PACT median CF is included for reference. Error bars represent median ± median absolute deviation.

**Extended Data Figure 11:**
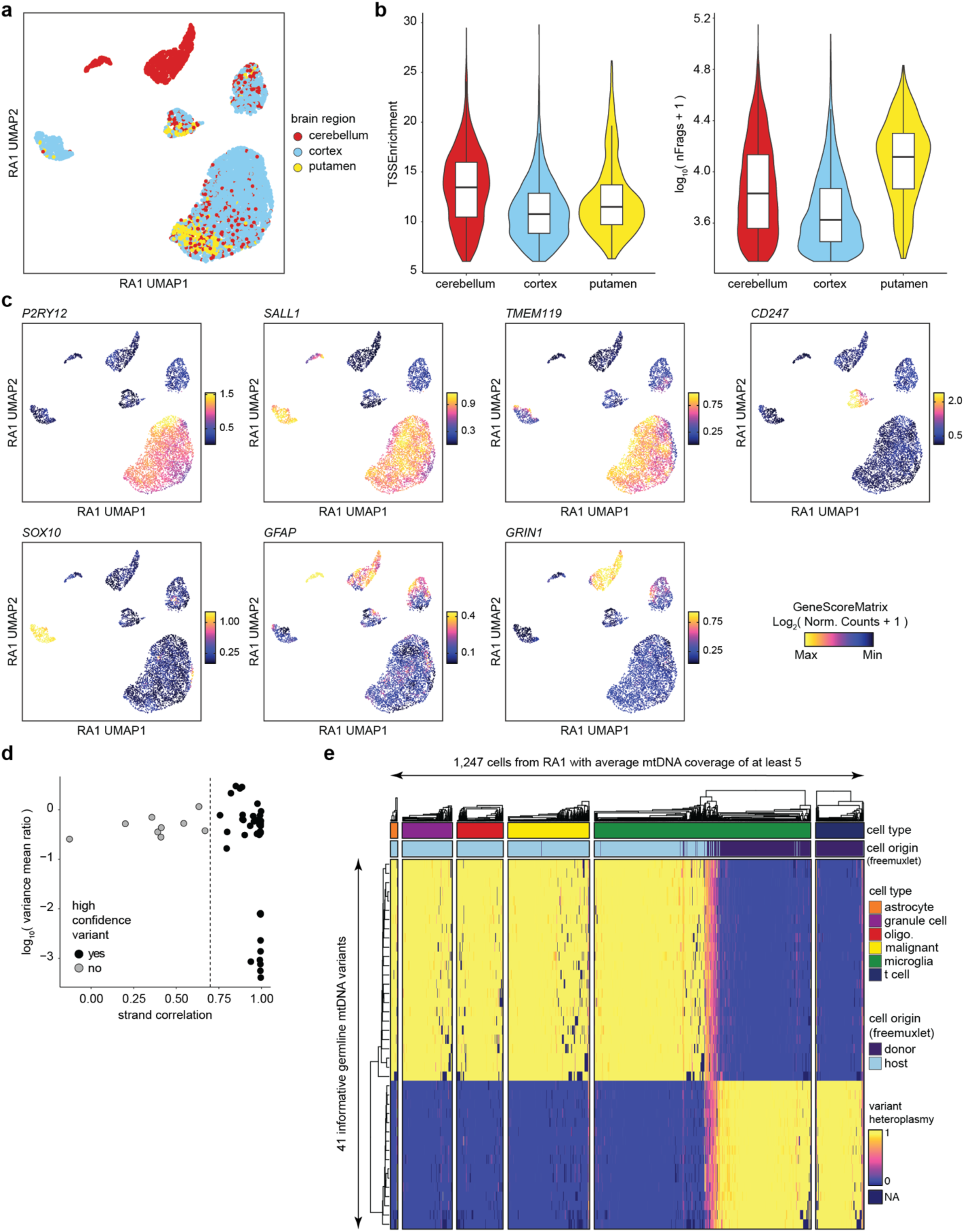
Additional mtscATAC-seq data from BMT recipient RA1. **(a)** scATAC-seq profiles of cells from RA1 colored by brain region. **(b)** TSS enrichment per cell (left) and number of fragments per cell (right) for each brain region profiled from RA1. **(c)** scATAC-seq profiles of cells from RA1 colored by gene accessibility for selected marker genes. **(d)** mtDNA variants identified by mgatk from aggregated RA1 data. **(e)** Concordance of mtDNA genotypes and germline genotypes in single cells from RA1 data. mtDNA genotypes were determined by mgatk; germline genotypes were determined by freemuxlet.

**Extended Data Figure 12:**
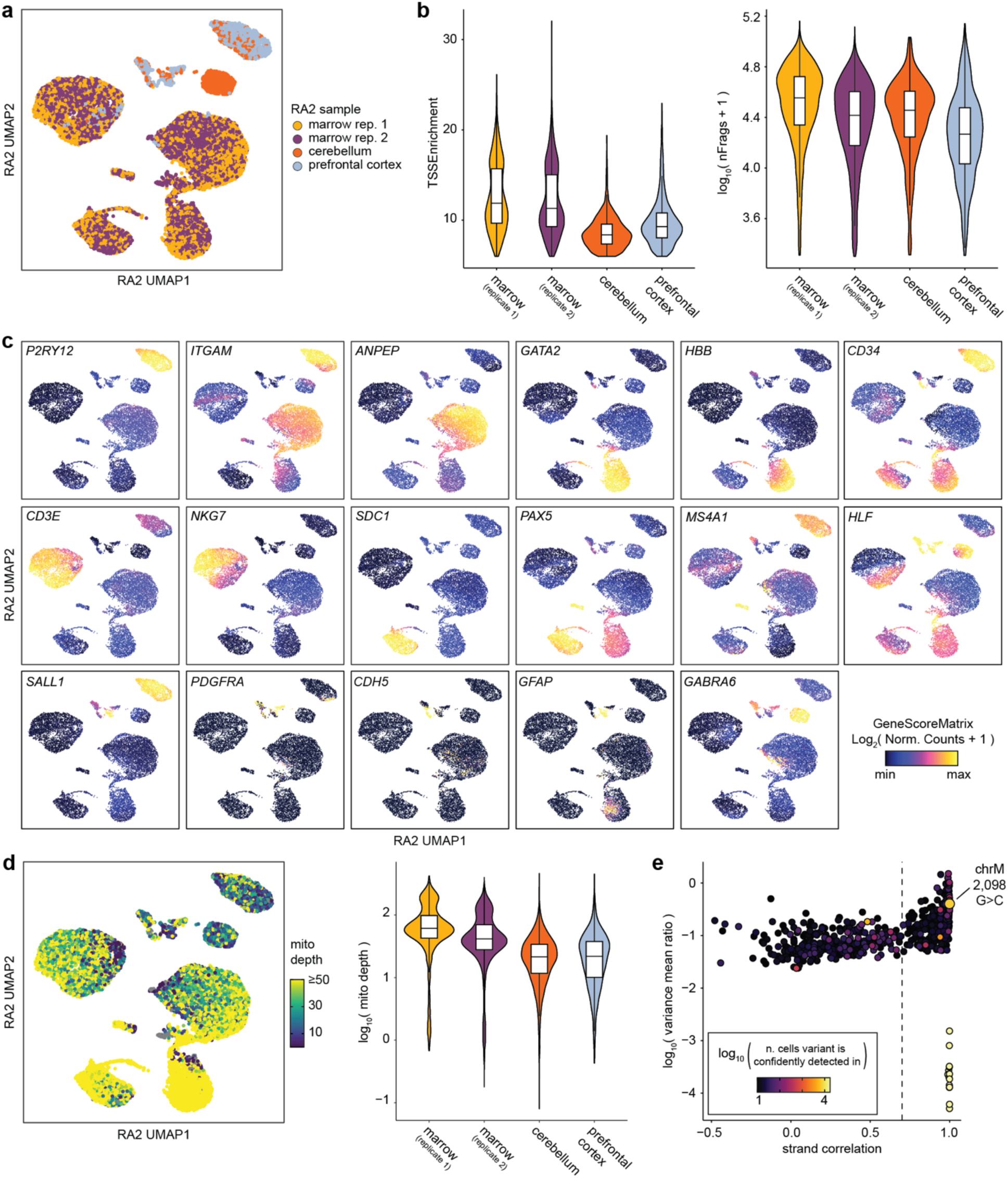
Additional mtscATAC-seq data from 77-year-old RA2. **(a)** scATAC-seq profiles of cells from RA2 colored by sample. **(b)** TSS enrichment per cell (left) and number of fragments per cell (right) for each sample profiled from RA2. **(c)** scATAC-seq profiles of cells from RA2 colored by gene accessibility for selected marker genes. **(d)** Mitochondrial genome coverage as quantified by mgatk in single cells (left) or aggregated by sample (right). **(e)** mtDNA variants identified by mgatk from aggregated RA2 data.

**Extended Data Figure 13:**
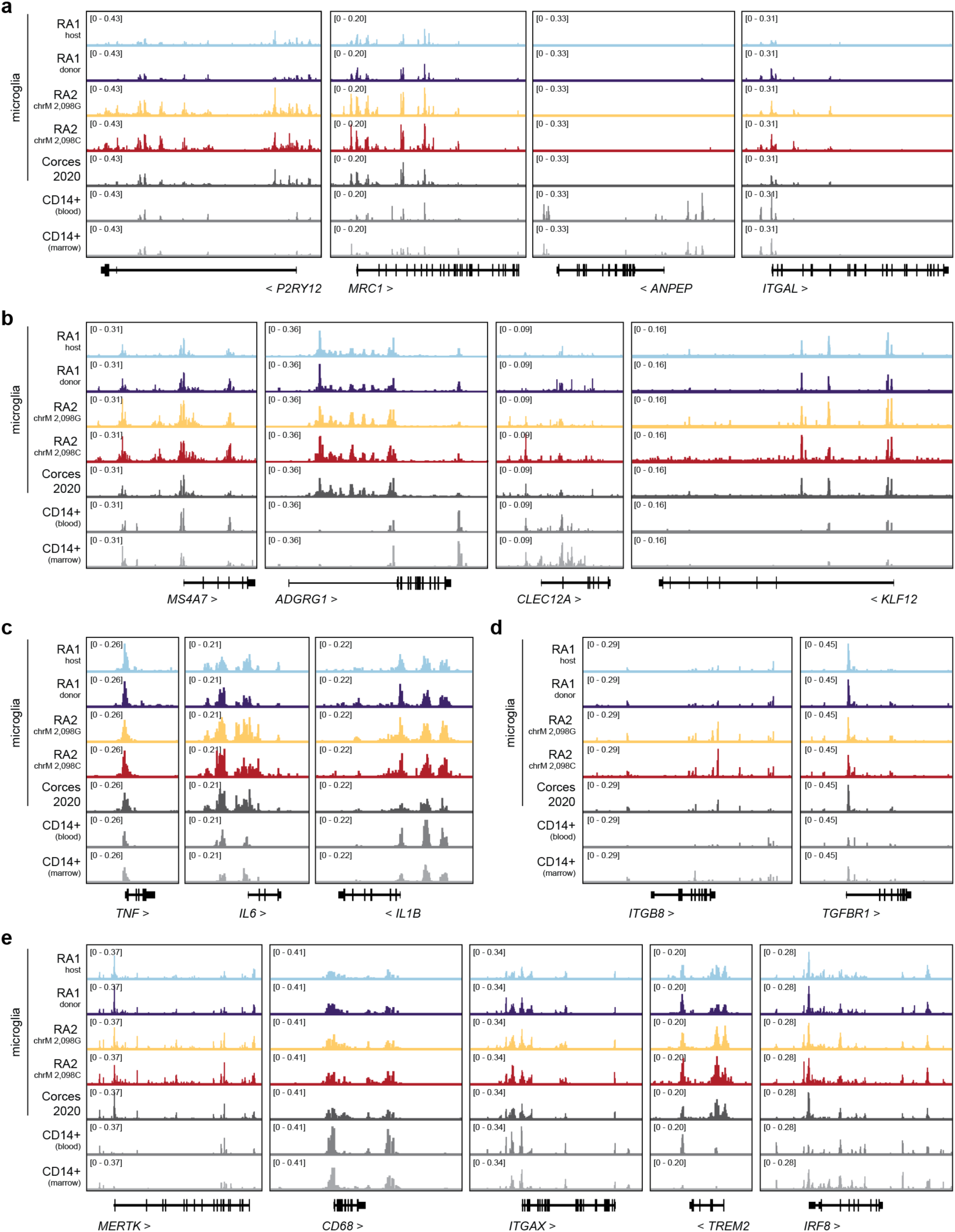
Infiltrating myeloid cells in the brain of RA1 and RA2 are near-identical to microglia at additional gene loci. Pseudobulk scATAC-seq tracks for **(a)** myeloid marker genes, **(b)** brain-infiltrating myeloid cell marker genes, **(c)** cytokines, **(d)** TGFβ-related genes, and **(e)** phagocytosis and disease-associated microglia genes.

**Extended Data Figure 14:**
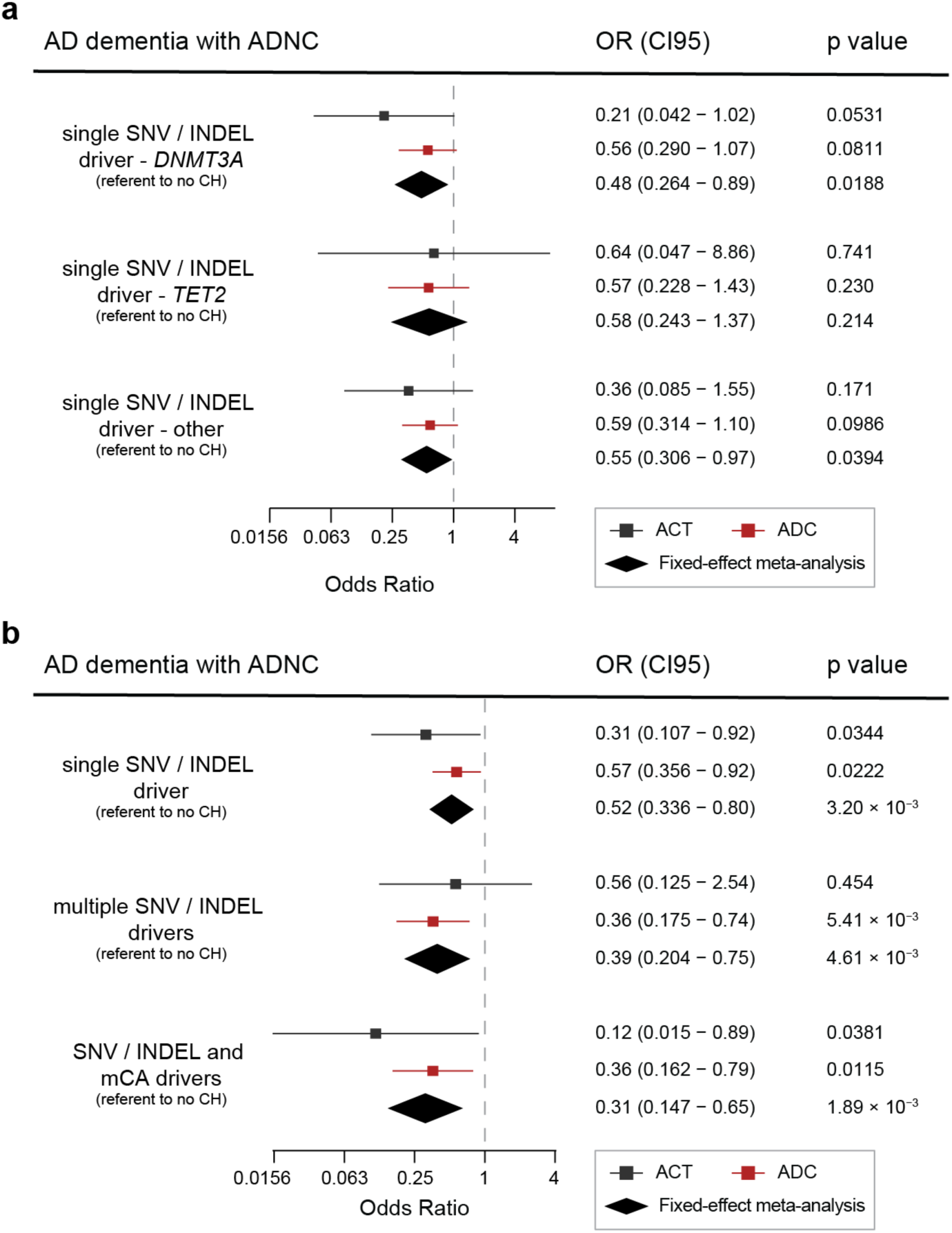
Associations of CH and AD by mutated gene and by number of drivers. **(a)** Effect of mutated gene on the risk of AD dementia with ADNC. **(b)** Effect of single and multi-driver CH on the risk of AD dementia with ADNC. **(a-b)** include n=192 subjects in the ACT cohort and n=1,273 subjects in the ADC cohort. Additional demographic information is available in Supplementary Tables 3 and 4. OR, CI95, and two-sided Wald P value were calculated from logistic regression models which included age, sex, and *APOE* genotype as covariates. The measure of center is the OR and the lengths of the lines represent the CI95 for the OR. For meta-analyses, the center of the diamonds represents the OR and the lengths represent the CI95 for the OR.

**Extended Data Figure 15:**
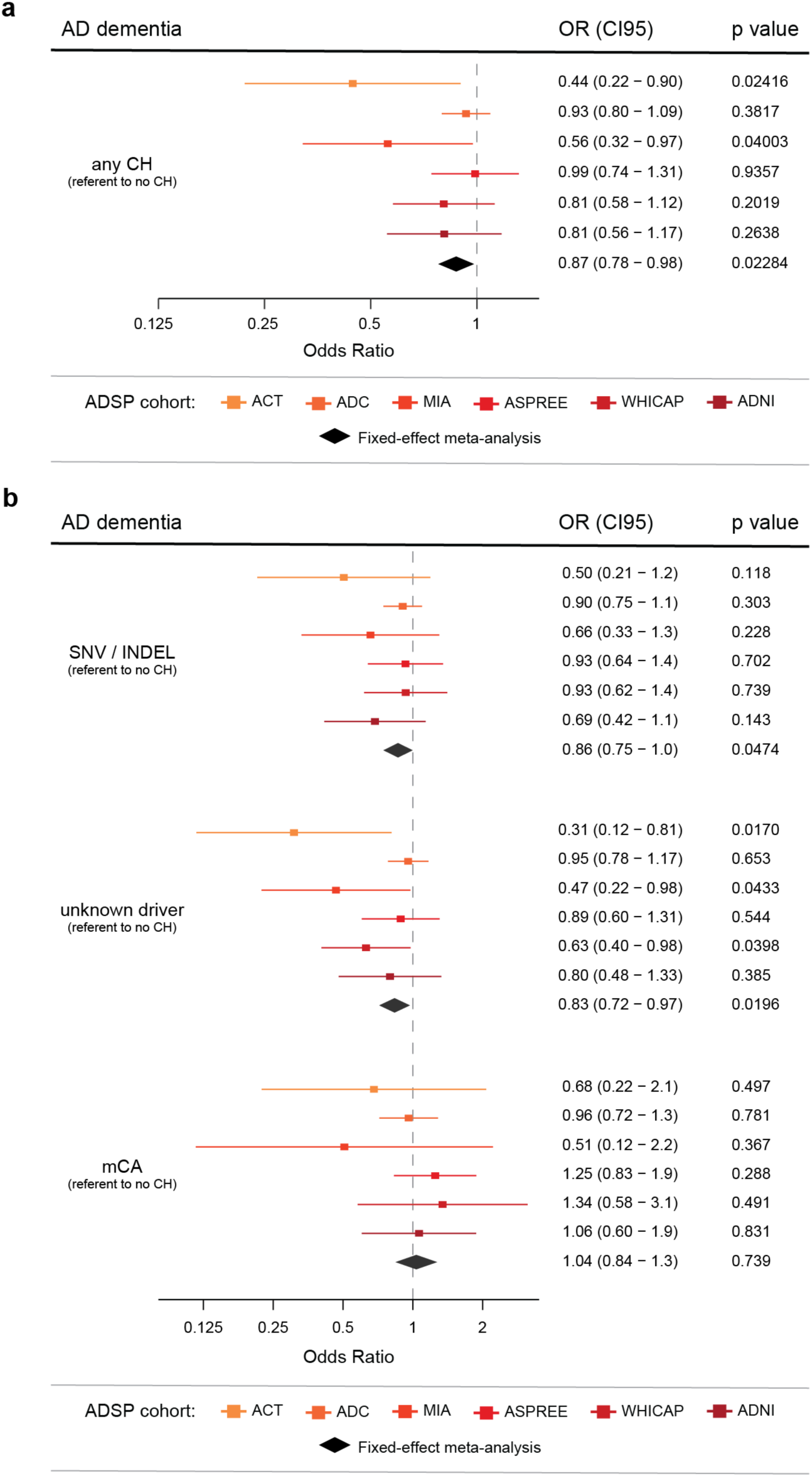
Associations of CH and AD dementia in additional ADSP cohorts. **(a)** Effect of CH on the risk of AD dementia. **(b)** Effect of CH driver type on the risk of AD dementia. **(a-b)** include individuals in the ACT (n=172), ADC (n=3,070), MIA (n=353), ASPREE (n=3,066), WHICAP (n=701), and ADNI (n=990) cohorts. Additional demographic information is available in Supplementary Table 15. For longitudinal cohorts (ASPREE, WHICAP, and ADNI) only incident AD cases were considered. For other cohorts, analysis was restricted to individuals older than 75 at blood draw and older than 65 at time of AD diagnosis. OR, CI95, and two-sided Wald P value were calculated from logistic regression models which included age, sex, and *APOE* genotype as covariates. The measure of center is the OR and the lengths of the lines represent the CI95 for the OR. For meta-analyses, the center of the diamonds represents the OR and the lengths represent the CI95 for the OR.

## Methods

### PACT – postmortem human brain specimens

Postmortem brain tissue and donor metadata were obtained via the UW BioRepository and Integrated Neuropathology (BRaIN) laboratory from participants in the Kaiser Permanente Washington Health Research Institute Adult Changes in Thought (ACT) Study and the University of Washington AD Research Center (ADRC). ACT is a longitudinal, community-based observational study of brain aging in participants older than 65 randomly sampled from the Group Health Cooperative (now Kaiser Permanente Washington), a health management organization in King County, Washington. A subset of participants in the study donate their brains for research upon death, and a comprehensive neuropathological exam is performed to assess for neurodegenerative disease pathologies including AD^58^. For decedents with postmortem intervals of less than 12h, a rapid autopsy is performed in which samples from multiple brain regions are taken from one hemisphere and flash-frozen in liquid nitrogen. Consent for brain donation was obtained from each donor and the study was approved by the University of Washington Institutional Review Board and by the Kaiser Permanente Washington Institutional Review Board. We obtained occipital cortex samples from ACT2-7 which we have previously investigated^5^ and additional subjects UW1-23 (**Supplementary Table 2**).

### PACT – design of hybridization capture panels

#### Variant calling and annotation

WGS of ADSP samples, including the ACT cohort, was previously performed^59^. Read data aligned to the hg38 reference genome was downloaded in CRAM format from Amazon Web Services using the National Institute on Aging Genetics of Alzheimer’s Disease (NIAGADS) Data Sharing Service^60^. GATK (v4.6.0.0) was used to process each CRAM file^61^. GATK Mutect2 was used to identify putative somatic variants using tumor-only mode with default settings^62^. Putative variants identified by Mutect2 were filtered to identify known CHIP variants (SNV/INDEL driver variants) as well as passenger variants, as previously described^5,63^. Briefly, to identify driver SNV/INDELs, we used a curated whitelist of variants in 73 genes (**Supplementary Table 1**). *U2AF1* mutations require a separate pipeline due to an erroneous duplication in the hg38 reference genome, so we used our previously reported strategy to identify these variants^63,64^.

To identify high-confidence passenger variants, we used our previously published approach^63^. Briefly, we excluded variants that failed one or more Mutect2 filters, required variants to have a sequencing depth between 20 and 100 (inclusive) and only considered biallelic single nucleotide variants. We removed variants present in low complexity regions of the genome or segmental duplications. To exclude germline contaminants, we removed any variant appearing in the TOPMed Freeze 8 callset (705 million variants from 132,354 whole genomes). To identify mosaic chromosomal alterations, we downloaded the germline variant calls for each individual in VCF format from the NIAGADS Data Sharing Service. We used the MoChA pipeline with default settings^65,66^. We only included mCA variants with strong evidence of involvement in myeloid malignancy (Loss of Y chromosome, 7q-, 5q-, 20q-, *DNMT3A* deletion or loss of heterozygosity [LOH], *TET2* deletion or LOH, *JAK2* LOH).

#### Design of hybridization capture panels

For the custom hybridization capture panels used for the PACT protocol, we sought to maximize the number of informative mutations and included all passengers with an allele frequency less than 0.40, without performing the singleton filtering or C>T T>C filtering steps used below in the human genetic analyses. Once mutation sites of interest were identified (including passengers and SNV/INDEL driver mutations of interest) custom panels were designed and synthesized as xGen Custom Hyb Panels from Integrated DNA Technologies (IDT). We typically merged the variants from 5-10 individuals into a single panel, since we found that panels could be cost-effectively synthesized from IDT with up to 10,000 probes.

### PACT – experimental workflow

#### Nuclei isolation

Nuclei were isolated from frozen human postmortem brain tissue following a modified sucrose gradient ultracentrifugation protocol. Briefly, 100–200 mg of frozen tissue (kept on dry ice until use) was thawed for ∼5 min in 5 ml lysis buffer [0.32 M sucrose, 5 mM CaCl₂, 3 mM Mg(acetate)₂, 0.1 mM EDTA, 10 mM Tris–HCl (pH 8), 1 mM DTT, and 0.1% Triton X-100 in nuclease-free H₂O] on ice. Tissue was homogenized in a pre-chilled glass douncer (∼30 strokes) and transferred to a 50 ml ultracentrifuge tube. The volume was brought to 12 ml with lysis buffer, and 21 ml sucrose buffer [1.8 M sucrose, 3 mM Mg(acetate)₂, 1 mM DTT, 10 mM Tris–HCl (pH 8)] was gently added from the bottom to form a density gradient. Tubes were balanced, placed in a SW32Ti swinging-bucket rotor (Beckman Coulter), and ultracentrifuged at 25,000 RPM (107,163 × g) for 2 h at 4 °C. The supernatant was discarded, and 500 μl of 1× PBS containing 1% BSA was added to the pellet. Pellets were incubated on ice for 10 min, resuspended, and transferred to 5 ml filter-cap tubes (12×75mm).

#### Nuclei staining and flow cytometry

Nuclei suspension was centrifuged at 500 × g for 5 min. Nuclei were resuspended at 200,000 cells in 50 μl of 0.5% BSA in 1× PBS, then stained for 30 min on a shaker with Anti-NeuN Alexa Fluor 700 (1:400) and Anti-Olig2 Alexa Fluor 488 (1:100). Nuclei were washed and sorted into Zymo genomic lysis buffer (data acquired with FACSDiva v8.0.1, BD Biosciences).

#### DNA extraction, library preparation, hybridization capture, and sequencing

Genomic DNA was extracted using the Quick-DNA Microprep Kit (Zymo Research) following the manufacturer’s instructions. Samples were lysed in Genomic Lysis Buffer (4:1 buffer:sample), bound to Zymo-Spin IC columns, washed with DNA Pre-Wash Buffer and g-DNA Wash Buffer, and eluted in 30 µl DNA Elution Buffer. Genomic DNA was fragmented, end-repaired, and dA-tailed using the Twist Library Preparation EF Kit 2.0 (Twist Bioscience). Twist Universal Adapters were ligated, and products were purified with 0.8× DNA purification beads (Twist). Libraries were PCR-amplified (10 cycles) with Twist UDI primers and Equinox Library AMP Mix, followed by 1× bead purification. Hybridization capture was performed using the xGen Hybridization and Wash Kit (Integrated DNA Technologies) with xGen Lockdown Probes. Libraries were hybridized at 65 °C for 16 h, captured with streptavidin-coated magnetic beads, washed at 65 °C, PCR-amplified (11 cycles), and purified with 1× DNA purification beads. Libraries were sequenced on an Illumina NovaSeq X Plus sequencer using paired end 150×150 base pair reads.

### PACT – analytical workflow

#### Quantification of passengers and SNV/INDEL drivers

Fastq files were trimmed with fastp using --detect_adapter_for_pe. Trimmed reads were aligned to the hg38 genome using bwa. Reads were filtered based on q30 values and then duplicates were removed using Picard. Passenger allele frequencies were assessed by computing read pileups using the samtools mpileup utility, using as input the de-duplicated bam file for each sample and a bed file containing the locations of all passenger mutations in the panel. If a particular sample contained known SNV/INDEL driver mutation(s) in the blood, these were also included in the bed file for quantification with samtools mpileup. For variant discovery, Mutect2 was used as described above for variant calling and putative variants were filtered using a curated whitelist.

#### Variant filtering and clone identification

Passenger mutations for each individual were filtered on coverage, requiring all samples to have a coverage of at least 200. Mutations with a CF < 0.002 were considered not detected. Finally, mutations were filtered based on having a myeloid-enriched CF at least 2x that of the total brain CF. Non-specific mutations that failed this filter likely represent germline variants or variants occurring very early in development. Finally, clones were assigned based on manual identification of modes in the myeloid-enriched CF distribution. Typically one clone was clearly identified, but in certain cases two or even three clones were apparent. Finally, occasionally outlier variants were detected that had a myeloid-enriched CF well above that of other variants. While these variants had minimal effect on the clone median, when present, these were labeled as “outliers” and were not included in the primary clone for visualization purposes.

### PACT – estimation of percent mutant microglia

#### snATAC-seq protocol

Nuclei isolation from frozen brain specimens was performed as described above. Nuclei suspensions were split into two samples, one which was used for snATAC-seq, and an identically-treated sample which was used for genomic DNA isolation and PACT mutation profiling as described above. Single nucleus ATAC-seq libraries were prepared using the Chromium Next GEM Single Cell ATAC Kit (10x Genomics Cat#1000390). Nuclei were transposed with Tn5 transposase, partitioned into Gel Bead-in-Emulsions (GEMs) for barcoding and amplification, purified with SPRIselect beads (Beckman Coulter), and indexed by PCR. Libraries were size selected, quantified, and assessed on an Agilent Bioanalyzer.

#### snATAC-seq pre-processing and clustering

snATAC-seq clustering was performed as described previously^5^. Read trimming, deduplication, and alignment was performed using the cellranger-atac count pipeline from 10X Genomics (v2.1.0) and the hg38 reference genome. Fragments files from cellranger-atac count into ArchR (v1.0.3) for downstream clustering and analysis^67^. Cell calling proceeded in two steps – first, barcodes that had a TSS enrichment of at least 4 and at least 2000 fragments were retained and initial processing and clustering was performed. Secondly, a cluster of debris was identified by low nFrags and low TSSEnrichment, excluded, and then the remaining nuclei were re-clustered. Processing proceeded according to the standard ArchR workflow, including creation of Arrow files, doublet simulation and removal, dimensionality reduction using addIterativeLSI, clustering, and visualization via UMAP. Harmony batch correction was not performed. To aid in cluster interpretation, previously published snATAC-seq datasets from the brain and blood were clustered together with our samples^35,36^.

#### Estimation of percentage of mutant microglia

We adapted our previously described procedure for estimating the percentage of mutant microglia to use the median CF of the PACT passenger variants^5^. We first determined the number of microglia and non-microglial hematopoietic cells in each brain sample. We conservatively assumed that all non-microglial hematopoietic cells were mutant, therefore we subtracted out the expected contribution of mutant alleles from these non-microglial cells to calculate an adjusted CF for each sample where mutant alleles could only be contributed by microglia. We then divided the adjusted percent mutant cells in each sample by the percent microglia in each sample to calculate the percentage of mutant microglia. To obtain a 95% confidence interval, we then estimated bootstrapped confidence intervals for percent mutant microglia for each sample using a binomial simulation based on input parameters for VAF, percent microglia, and number of cells unique to each sample.

#### Detection of the Y chromosome in snATAC-seq data

We sought to classify individual nuclei as LoY or WT from our snATAC-seq data by counting reads mapping to the Y chromosome in each nucleus. For this analysis, we investigated snATAC-seq data from individuals with LoY present in their blood (UW4; UW12-UW16) and restricted our analysis to the 26,435 adequately sequenced nuclei from these individuals (nFrags ≥ 10,000). To remove artefactual reads mapping to the Y chromosome, we took the 10X cellranger-atac peak calls from the Y chromosome from a female sample. We expanded each of these peaks 10kb upstream and downstream to create a blacklist. We then considered the Y chromosome fragments for UW4 and UW12-UW16 and removed any fragments overlapping our chrY blacklist. Finally, we counted the number of chrY fragments in each nucleus and added this information to the ArchR project metadata. Individual nuclei were called as LoY if they had no more than 2 ATAC-seq fragments mapping to the Y chromosome.

### mtDNA lineage tracing – fresh human tissue specimens

Postmortem tissues specimens were obtained through Stanford Research Autopsy Center at Stanford University School of Medicine. Informed consent for research autopsy was obtained by the Coordinator of the Research Autopsy Center at Stanford and reviewed at the time of death prior to tissue collection. All procedures were approved by the Institutional Review Board (IRB) of Stanford University School of Medicine (IRB#63818) and conducted in accordance with relevant ethical guidelines. At autopsy, the brain was removed, bisected, coronally sectioned, and samples from indicated brain regions were collected. Lumbar vertebrae were removed and separated. Brain tissues and vertebra were immediately placed in ice-cold RPMI 1640 medium (Thermo Fisher Cat#11875093). The warm postmortem interval was 2 hours 26 minutes for RA1 and 2 hours 8 minutes for RA2.

### mtDNA lineage tracing – experimental workflow

#### Single cell isolation from fresh human brain specimens

Each brain region was cut into 2g pieces, minced, and placed into individual GenteMACS C tubes. Enzymatic digest was performed according to the Neural Tissue Dissociation Kit (P) (Cat# 130-092-628; Miltenyi Biotec). Briefly, an enzyme cocktail consisting of 250µl Enzyme P, 50µl Enzyme A, 9500µl Buffer X, 100µl Buffer Y was added to each tube. The mixture was run through a GentleMACS Octo Dissociator (37C_NTDK_1) for 22 minutes. 10ml HBSS (with Calcium and Magnesium) was added to the suspension and filtered through a 100µm filter into a new 50ml tube, and an additional 10ml of HBSS (with Calcium and Magnesium) was used to rinse the filter. Each sample was centrifuged at 300 x g for 10 min at 4 °C. Upon removing the supernatant, the pellet was resuspended with 12.4 ml ice-cold PBS. 3.6 mL Debris Removal Solution was then added and mixed well. Finally, 8mL ice-cold PBS was overlayed on top of the mixture. Each gradient was centrifuged at 3000x g for 10min at 4 °C. The supernatant containing the removed myelin debris was aspirated. The remaining pellets were resuspended and pooled with a total volume of 3mL of 1x RBC Lysis and incubated for 5min. The RBC Lysis was quenched with 7mL PBS and centrifuged at 300 x g for 5 min at 4 °C. The supernatant was aspirated, and the pellet was resuspended with 1mL of PBS for staining.

#### Bone marrow isolation from human vertebral bodies

To isolate bone marrow cells from, an individual vertebral body was stabilized onto a clamp. The white intervertebral disk was carefully removed to expose the bone marrow. The bone marrow was scooped out of the vertebral body and placed in a mortar containing 10mL HBSS (with Calcium and Magnesium). Using a pestle, the bone marrow was gently crushed for 1min. Liquid was removed and filtered through a 100µm filter into a new 50mL tube. Cells were centrifuged at 300 x g for 10 min at 4 °C. The supernatant was aspirated, and the pellet was resuspended with 10mL 1x RBC Lysis for 5-7min. The sample was quenched with 30mL PBS, and centrifuged 300 x g for 5 min at 4 °C. The supernatant was aspirated and resuspended in 10mL of PBS for staining.

#### Staining and sorting of fresh human tissue specimens

Each sample was centrifuged at 300 x g for 5 min at 4 °C. The supernatant was aspirated, and up to 2 million cells were resuspended with 300µl FACS buffer (PBS with 2% FBS). The cells were then transferred through a filtered FACS tube on ice. 6µl FC Block (1:50), was added and placed on a shaker for 30 minutes. On ice, 0.3µl Fixable Viability Stain 700 (1:1000), 3µl CD45 Brilliant Violet 605 (1:100), 3µl CD66b APC (1:100), 3µl CD11b PE-Cy7 (1:100), and 0.3µl CalceinAM FITC (1:1000) were added to the sample, covered from light, and placed on a shaker for 45 minutes. Cells were washed twice. Live CD66b- cells were isolated via flow cytometry.

#### mtscATAC-seq protocol

Cells were pelleted at 500 × g for 5 min at 4 °C, resuspended in PBS, and fixed by adding 16% formaldehyde to a final concentration of 1% for 10 min at room temperature with occasional mixing. The reaction was quenched with 2 M glycine (0.125 M final), followed by centrifugation and washes with ice-cold PBS and wash buffer. Fixed cells were lysed in chilled lysis buffer, washed, and counted using trypan blue staining. Cell concentrations were adjusted according to the 10x Genomics recommendations, and samples were processed using the Chromium Single Cell ATAC workflow for transposition, barcoding, and library preparation.

### mtDNA lineage tracing – analytical workflow

#### mtscATAC-seq pre-processing, clustering, and differential analysis

Read trimming, deduplication, and alignment was performed using the cellranger-atac count pipeline from 10X Genomics (v2.1.0). We used a previously described custom hg38 reference genome which has been hard masked to prevent spurious alignments of mtDNA to the nuclear genome, which is standard practice for mtDNA variant calling^28,38^. We then imported fragments files from cellranger-atac count into ArchR (v1.0.3) for downstream clustering and analysis^67^. Barcodes that had a TSS enrichment of at least 6 and at least 2000 fragments were retained as cells for additional analysis. Processing proceeded according to the standard ArchR workflow, including creation of Arrow files, doublet simulation and removal, dimensionality reduction using addIterativeLSI, clustering, and visualization via UMAP. Harmony batch correction was not performed.

#### mtDNA variant calling

mtDNA variant calling was performed using mgatk v0.6.6^38^. To ensure uniform variant calls across all cells and all samples from a particular donor, bam files from each sample were merged into a single aggregated bam file per donor prior to variant calling. Mitochondrial reads from each sample were extracted into a separate bam file using samtools view^68^, and cell barcodes were renamed by substituting the standard “-1” barcode suffix with a custom suffix (e.g., −2, −3, etc) to preserve barcode uniqueness. Then all chrM bam files were merged using samtools merge^68^. Mgatk was run in tenx mode on the aggregated bam file. High confidence variants were identified using a strand correlation of at least 0.7 and then were added to the ArchR project metadata for additional visualization and analysis.

#### Genetic demultiplexing of donor and host cells within RA1

Freemuxlet (implemented in popscle v0.1) was used to deconvolve the donor and host genotypes within cells from bone marrow transplant recipient RA1^39,40^. To obtain a set of common variants to use for demultiplexing, we variants from the 1000 Genomes project available at http://ftp.1000genomes.ebi.ac.uk/vol1/ftp/data_collections/1000_genomes_project/release/20181203_biallelic_SNV/ALL.wgs.shapeit2_integrated_v1a.GRCh38.20181129.sites.vcf.gz. We then filtered these variants to retain only variants with an allele frequency of at least 0.01 and excluded variants present in the ENCODE blacklist. We removed duplicates from the mtscATAC-seq bam files using picard markDuplicates, computed pileups of each bam file at each variant of interest using popscle dsc-pileup and then ran freemuxlet using popscle freemuxlet with --nsample set to 2. The SNG.BEST.GUESS column of the freemuxlet output was used to assign genotypes to each cell. Freemuxlet 0 or 1 output values were manually recoded to “donor” based on which genotype was prevalent in the T cell cluster (expected to be mostly donor cells) or “host” based on which genotype was prevalent in the non-hematopoietic brain cell types (expected to be host cells). Freemuxlet genotypes were further confirmed using germline mtDNA mutations.

### Human cohort data – study populations and variant calling

#### ADSP study populations

The ADSP is a collaborative effort of the National Institutes of Aging, the NHGRI and the Alzheimer’s community to understand the genetic basis of AD^59^. In release 4 of the ADSP WGS dataset, WGS data are available from 35,014 participants across 17 cohorts^69^. Cases met NINCDS–ADRDA criteria for possible, probable or definite AD, had documented age at onset or age at death, and APOE genotyping. Written informed consent was obtained from all human participants in each of the studies that contributed to ADSP in our analysis, which are listed below. Secondary analysis of the dbGaP data in this manuscript was approved by the Partners Healthcare and Stanford University Institutional Review Boards, and this work is compliant with all relevant ethical regulations.

Additional information about each cohort is available at:

- ACT^58^ – see above (**PACT – postmortem human brain specimens**) and https://dss.niagads.org/cohorts/adult-changes-in-thought-act/
- ADC – https://dss.niagads.org/cohorts/nia-alzheimers-disease-research-centers-adrc/
- MIA – https://dss.niagads.org/cohorts/university-of-miami-mia/
- ASPREE^70^ – https://dss.niagads.org/cohorts/asprin-in-reducing-events-in-the-elderly-aspree/
- WHICAP^71^ – https://dss.niagads.org/cohorts/washington-heights-and-inwood-community-aging-project-whicap/
- ADNI^72^ – https://dss.niagads.org/cohorts/alzheimers-disease-neuroimaging-initiative-adni/

#### Variant calling and clonal hematopoiesis subtype identification

Driver events were defined as described above for both SNV/INDEL and mCAs. We used cell fraction minimum of 0.08 for mCA drivers. We did not use a minimum VAF threshold for SNV/INDEL drivers, because WGS is already limited in its ability to detect small clones. Individuals with both SNV/INDEL and mCA drivers were included in the “SNV/INDEL” category. To define CH with unknown drivers, we relied on the principal that only expanded clones should have detectable passenger mutations in low depth sequencing assays. First, we performed additional filtering beyond the filtering performed for the PACT custom panels, in order to identify a higher-confidence set of passenger variants. Specifically, we included only C>T and T>C base substitutions and performed singleton filtering, removing any variant that appeared more than once across all processed samples. Finally, we removed any variant with an AF larger than 0.30.

We sought to empirically identify a threshold passenger mutation count above which we had high confidence that a clone was present. Age and two germline variants in *TCL1A* (rs2887399) and *TERT* (rs7705526) are known to associate strongly with CH^73^, so we identified an optimal threshold for categorizing each sample as CH with unknown driver or no CH based on the strength of the association of presence of CH with these three variables using logistic regression with sex, self-reported race, and cohort as additional covariates. All six ADSP cohorts (ACT, ADC, MIA, ASPREE, WHICAP, and ADNI) that are described below were used for this analysis. We selected different thresholds for testing and found that having greater than negative 0.5 standard deviations (30.8th percentile) of the mean passenger count in those with CH with known drivers was optimal based on ranking the p-values for age, rs2887399 [*TCL1A*], and rs7705526 [*TERT*]. Individuals above this threshold of passenger variants without known drivers were considered to have CH with unknown drivers. The percentile threshold was applied for each cohort withing ADSP separately due to variability from cohort to cohort in the total number of passenger mutations identified by WGS.

### Human cohort data – statistical analysis plan

#### Cohorts with postmortem neuropathology

To test for associations between all forms of clonal hematopoiesis with AD, we used data from ACT and ADC cohorts within the ADSP that had both dementia diagnoses as well as postmortem neuropathological characterization. We classified participants as having either low (Braak stage 0-3 and CERAD score 0-1 OR Braak stage 0-2 and CERAD score 2), high (Braak stage 5-6 and CERAD score 2-3), or intermediate (all others) ADNC. Neuropathologically confirmed AD was defined as having intermediate/high ADNC and clinical dementia. We then performed logistic regression package in R with neuropathologically confirmed AD as the outcome variable and age at death, sex, APOE genotype, and clonal hematopoiesis or clonal hematopoiesis subtypes as input variables. Results from ACT and ADC were analyzed separately and the results were meta-analyzed using a fixed effects model from the meta package in R.

#### Additional ADSP cohorts

We sought to replicate the findings from the neuropathologically-confirmed cohorts in additional cohorts with only clinical dementia diagnoses. For this purpose, we used subjects enrolled in ACT, ADC, MIA, ASPREE, WHICAP, and ADNI, which are also part of the ADSP. Due to privacy concerns, those age 90 or older did not have an exact age available on dbGaP, and were considered to be age 90 for the purposes of this analysis. For all cohorts, separate statistical associations were performed using CH as a two-factor variable (any CH versus no CH) and as a four-factor variable (SNV/INDEL-driven CH, mCA-driven CH, or CH with unknown drivers versus no CH).

For case-control cohorts ACT, ADC, and MIA, where assessment of CH from blood DNA was often done several years after AD dementia diagnosis, we excluded individuals who were younger than 75 at study enrollment or developed AD younger than 65. These exclusions remove individuals who likely developed AD or had pre-clinical AD pathophysiology prior to the development of a CH clone of appreciable size. For each case-control cohort, we used logistic regression to test for an association between CH and AD. Age at blood draw, sex, and *APOE* genotype were included as covariates. For longitudinal cohorts ASPREE, WHICAP and MIA, we restricted our analysis to incident AD cases and again used logistic regression to test for an association between CH and AD. Age at baseline, sex, and *APOE* genotype were included as covariates. A fixed-effect meta-analysis for the effect of CH subtypes on risk of AD in all six cohorts was then performed using the logistic regression results for each cohort in the R package meta.

## Data Availability

Whole genome sequencing data and phenotype data is available from the ADSP via NIAGADS for investigators with approved protocols^60^. Sequencing data generated in this study will be deposited on dbGaP and eligible investigators will be able to apply for access via the dbGaP Authorized Access system.

## Code Availability

Passenger variant identification: https://github.com/weinstockj/passenger_count_variant_calling

snATAC-seq analysis: https://github.com/juliabelk/CHIP_and_AD

mgatk: https://github.com/caleblareau/mgatk

## Acknowledgements

We thank Dr. Caleb Lareau and members of the Jaiswal and Chang labs for helpful discussions. This work was supported by grants from the Ludwig Cancer Center Research at Stanford University, Burroughs Wellcome Fund Career Award for Medical Scientists, Phil and Penny Knight Initiative for Brain Resilience at Stanford University, the National Institutes of Health (DP2-HL157540, 1R01AG088656, as well as 1R01AG088657) (S.J). J.A.B. is supported by an HHMI Hanna Gray Fellowship. H.Y.C. was an Investigator of the Howard Hughes Medical Institute. Cell sorting for this project was done on instruments in the Stanford Shared FACS Facility (RRID: SCR_017788). The UW BioRepository and Integrated Neuropathology (BRaIN) Laboratory and Precision Neuropathology Core are supported by the NIH via the UW Alzheimer’s Disease Research Center (P30 AG066509), the ACT study (U19 AG066567), the BRAIN Initiative Cell Atlas Network (UM1 MH130981 and UM1MH134812), the Seattle Alzheimer’s Disease Brain Cell Atlas (U19AG060909), multiple collaborative projects (U24AG072458; U24NS133949; U24NS133945; U24NS135651; U01NS137500; and U01NS137484), the US Department of Defense (DoD W81XWH-21-S-TBIPH2) and the Allen Institute for Brain Science. C.D.K is supported as the Nancy and Buster Alvord Endowed Chair in Neuropathology and as a Weill Neurohub Investigator. We thank Emily Ragaglia and Aimee Schantz for incredible administrative support, John Campos and Mark Montine for data management, and Jenna Kelley, Julia Ryan, Flavia Ernau, Kim Howard, and Katie Miller for outstanding technical support. Finally, we are deeply grateful to the research participants and their families without whom this work would be impossible.

## Author Contributions

J.A.B. and S.J. conceptualized the project. J.A.B. conceptualized the experimental strategy. J.A.B., H.Y.C., and S.J. supervised experiments. J.A.B., Y.Z., Q.S., L.M., R.K., A.M.E., E.E.R., J.W., R.L., A.E.E., N.W., D.P., A.C., M.R.C., and K.S. performed experiments. J.A.B. analyzed data. J.A.B., E.E.R., S. Bukhari, I.C., D.E.B., H.V., I.L.W., T.J.M., J.E.H., and C.D.K. provided access to human tissue specimens. A.C. and D.B. provided expertise on human brain nuclei experiments. A.E.E., N.W., S. Brioschi, J.G., I.T., A.B., S.M., M.C. and I.L.W. provided expertise on processing fresh human tissue specimens. S.R., D.R., S. Brioschi, M.C. and A.B. provided expertise on mtscATAC-seq. J.A.B. and S.J. performed human genetic association tests. D.C.N., C.M.A., and B.N.V. provided expertise on human genetic analyses. S.J. and H.Y.C. provided resources. J.A.B. wrote the manuscript with input from all authors.

